# Radiating despite a lack of character: closely related, morphologically similar, co-occurring honeyeaters have diverged ecologically

**DOI:** 10.1101/034389

**Authors:** Eliot T. Miller, Sarah K. Wagner, Luke J. Harmon, Robert E. Ricklefs

## Abstract

The 75 species of Australian honeyeaters (Meliphagidae) are morphologically and ecologically diverse, with species feeding on nectar, insects, fruit, and other resources. We investigated ecomorphology and community structure of honeyeaters across Australia. First, we asked to what degree morphology and ecology (foraging behavior) are concordant. Second, we estimated rates of trait evolution. Third, we compared phylogenetic and trait community structure across the broad environmental gradients of continental Australia. We found that morphology explained 37% of the variance in ecology (and 62% vice versa), and that recovered multivariate ecomorphological relationships incorporated well-known bivariate relationships. Clades of large-bodied species exhibited elevated rates of morphological trait evolution, while members of *Melithreptus* showed slightly faster rates of ecological trait evolution. Finally, ecological trait diversity did not decline in parallel with phylogenetic diversity along a gradient of decreasing precipitation. We employ a new method (trait fields) and extend another (phylogenetic fields) to show that while species from phylogenetically clustered assemblages co-occur with morphologically similar species, these species are as varied in foraging behavior as those from more diverse assemblages. Thus, although closely related, these arid-adapted species have diverged in ecological space to a similar degree as their mesic counterparts, perhaps mediated by competition.

## INTRODUCTION

Birds fly, whales swim, cheetahs run, and amoebae move (not very fast) via amoeboid movement. In organismal biology, the connection between form and function is intuitive and generally presumed to exist. When examined quantitatively, this assumption is often well supported (Miles and Ricklefs 1984; Miles et al. 1987; Saunders and Barclay 1992; Ricklefs and Miles 1994; Fitzpatrick et al. 2004; Leisler and Schulze-Hagen 2011). That species’ traits influence their local abundance is a central tenet of evolutionary biology (Darwin 1859; Tilman 1988; McGill et al. 2006). The connection between a species’ morphology, performance, and resource use—its ecology—has often led to the use of morphology as a surrogate for ecology. While ecology itself can be difficult to measure and variable across time (Lovette and Holmes 1995) and space (Suryan et al. 2000), morphology can be measured quickly and is assumed to integrate over the life span of the individual and to reflect past selective pressures. Morphological variation influences ecological relationships and determines where individuals of a species can survive and which species can coexist in local assemblages.

Of course, similar ecologies can be realized through dissimilar morphologies, and a single morphology can serve varied purposes (Wainwright 2007). Caterpillars and giraffes both feed on leaves, and while many kingfishers specialize on fish, others focus almost entirely on terrestrial prey. Evolutionary constraints at multiple levels shape how species do and do not adapt to their environment (Arnold 1992; Futuyma 2010). For instance, Allen’s rule has been shown to influence morphology in birds, with species from colder climes characterized by shorter beaks (Allen 1877; Symonds and Tattersall 2010), and perhaps shorter legs. Thus, beak size may not be free to evolve towards a given adaptive peak, as it may be subject to conflicting selective pressures and, if beak length and leg length are genetically correlated, then evolution in some morphological directions may be constrained (Schluter 1996).

A species’ evolutionary history—the past selective regimes that have shaped its gene pool—may also constrain its adaptations. The ancestral state of a population can also influence which local adaptive peak it settles on (Hansen and Houle 2008). In addition, a given resource might be exploited by organisms sitting on several adaptive peaks, as in the case of caterpillars and giraffes feeding on the same leaves. Thus, while we expect some relationship between morphology and ecology among phylogenetically restricted groups of species, some authors have concluded that at a broader level “the strong influence of phylogeny within the trophic relationships of an assemblage negate[s] the value of an ecomorphological analysis” (Douglas and Matthews 1992). The amount of variance that morphology can explain in ecology, and whether tight ecomorphological relationships exist across large clades, warrants further study.

Environmental pressures push species within local assemblages to resemble each other, either through evolutionary processes or habitat filtering. Relatedly, if some lineages fail to colonize novel habitats from their ancestors, then across large climatic gradients these environmental pressures can lead to a reduction in phylogenetic diversity (Latham and Ricklefs 1993; Wiens and Donoghue 2004), presumably associated with a concomitant loss in morphological diversity. Yet, cursory examination of any naturally co-occurring set of species emphasizes that additional forces, particularly interspecific competition, act to reduce similarity among related species. This spectrum of possible community assembly processes forms the basis for studies on phylogenetic community structure (Webb et al. 2002). This proxy approach uses evolutionary relationships to indicate ecological similarity. A complementary approach to studying community assembly makes use of functional traits (McGill et al. 2006). Based largely on broad latitudinal trends in functional traits (Wright et al. 2004) and more limited ecophysiological assessments (Lambers and Poorter 2004), some authors have recently assessed the functional trait composition of local plant assemblages around the globe (Cornwell et al. 2006; Kraft and Ackerly 2010). Such studies are based on the assumption that these morphological traits have real-life ecological consequences within communities, and not just across communities. A functional trait approach based on morphology is itself a proxy for species’ ecologies.

Compared to the growing botanical literature, few modern studies have employed functional trait approaches to characterize bird assemblages (Luther 2009; Gómez et al. 2010; Ricklefs 2011; Ricklefs 2012; Dehling et al. 2014; Tobias et al. 2014), though this previously was an active area of research (Karr and James 1975; Ricklefs and Travis 1980; Keast and Recher 1997). Few studies have focused on ecological measures (e.g., foraging behavior), and little attention has been paid to changes in community organization across ecological gradients.

Limited phylogenetic diversity coupled with ecological opportunity may initiate a phase of adaptive radiation (Schluter 2000). While authors have differed on what constitutes an adaptive radiation (Givnish and Sytsma 2000; Schluter 2000; Givnish 2015), the general phenomenon of evolutionary diversification given ecological opportunity might characterize many lineages not traditionally recognized as being adaptive radiations. In such lineages we might expect to see increased rates of trait evolution compared to related lineages that have diversified in more stable environments. Thus, environmental pressures in combination with phylogenetic niche conservatism might lead to a loss of phylogenetic diversity, followed by a subsequent radiation in ecomorphological space by the lineages able to colonize these new areas.

The Australian honeyeaters (Meliphagidae) comprise a group of 75 species of passerine birds distributed across the continent, with at least one species found in almost every habitat type. Honeyeater species vary from large-bodied generalists like the Yellow Wattlebird (*Anthochaera paradoxa*, > 160 g) to small, decurved-billed nectarivores like the Red-headed Myzomela (*Myzomela* 7–8 g), and from stout-billed, ground-foraging insectivores like the Gibberbird (*Ashbyia lovensis*), to frequent frugivores like the Painted Honeyeater (*Grantiella picta*) (Higgins et al. 2001). A strong pattern of increasing phylogenetic clustering follows the gradient of decreasing precipitation, as mesic-adapted lineages drop out towards the arid interior of the continent (Miller et al. 2013). Yet, Meliphagidae species richness does not decline precipitously along this precipitation gradient, a fact that Miller et al. (2013) attributed, in part, to ecological opportunity and strong selective pressure to adapt to the newly opened desert habitats as Australia underwent dramatic aridification from the Miocene onwards.

In this paper, we address how evolution and ecology interact to determine species composition, local abundance, and trait diversity in honeyeater assemblages in Australia. We explore how these relationships change over climate gradients from both morphological and ecological perspectives. We ask whether trait diversity varies in parallel with phylogenetic diversity across local assemblages. When we find that it does not, we consider whether the phylogenetically limited subset of arid-adapted lineages has diversified phenotypically to fill trait space similarly to their mesic counterparts. Specifically, we test whether desert species evolved through trait space more rapidly, perhaps to fill ecological space left vacant as some lineages drop out along the precipitation gradient. We do so by analyzing two large, near-comprehensive datasets summarizing the morphological and ecological diversity of the Australian Meliphagidae. We predict the existence of strong ecomorphological relationships across the family (crown age ~ 20 mya), reflecting the biological axiom that form reflects function. Using the intersection of ecomorphology and distribution within the Australian continent (Miller et al. 2013), we test the prediction that lineages of phylogenetically clustered arid zone birds evolved rapidly in trait space to fill novel desert habitats and, accordingly, break the link between trait disparity and phylogenetic diversity.

## METHODS

### Morphological data collection and processing

We used digital calipers and photograph analysis (ImageJ, Schneider et al. 2012) to assemble a set of linear measurements on museum specimens: culmen length from front of the nares to bill tip; culmen length from base (kinetic hinge) of the bill to tip; exposed maxilla and bill chord (ImageJ); bill width and depth at both the nares and the base; wing chord (length from carpal joint to longest primary wing feather); length of the longest secondary wing feather; tarsus, hind toe and mid toe lengths; tail and total body lengths. Whenever possible, we measured at least 3 males and 3 females of each species/subspecies. We used ImageJ and spread-wing specimens to quantify total wing area, the length (along the axis of the wing) and width (widest point perpendicular to the wing axis) of the spread wing, and the lengths of the longest primary, the longest secondary, and the outermost/first secondary feather. We used these wing measurements to calculate the hand-wing index (supplementary material), which is a proxy for a wing’s aspect ratio, i.e., a measure of its shape, from rounded to pointed, generally associated with maneuverability versus long-distance flight and strong dispersal (Claramunt et al. 2012). Linear measurements are illustrated in the supplementary material.

We calculated the following additional indices: bill curvature, the quotient of the maxilla length over its chord (Rico-Guevara and Araya-Salas 2014); and a bill length index: 100× (bill length from base – bill length from nares)/bill length from base. Large values of the bill length index correspond to species where most of the length of the bill is proximal to the nares (e.g., Yellow Wattlebird, *Anthochaera paradoxa*), while small values correspond to species where most of the length is distal to the nares (e.g., Eastern Spinebill, *Acanthorhynchus tenuirostris*). We calculated bill width and depth indices in a similar fashion (using the width/depth at the base vs. the nares). Here, large values correspond to bills that taper considerably from base to tip in their width (White-gaped Honeyeater, *Stomiopera unicolor*) or their depth (Orange Chat, *Epthianura aurifrons*). These indices provide some indication of bill shape; many Tyrannidae flycatchers, for instance, have bills that are wide near the base and taper considerably towards the tip.

When available, we used the mass of the bird at collection as recorded on the specimen tag (37% of specimens). When no value was provided, we used the best information available to assign an approximate mass to each specimen. Specifically, if the sex of the specimen was known and a large sample of sex-specific, subspecies-specific masses was available (Higgins et al. 2001), we used that mass (59% of specimens). Otherwise, we used the next higher level of specificity, e.g., the sex-specific average mass across all subspecies of that species, etc. In this way, we assigned a mass to all specimens. We were unable to measure any specimens of the range-restricted Eungella Honeyeater (*Bolemoreus hindwoodi*), and excluded it from morphological analyses.

### Ecological data collection and processing

Collection of ecological information, primarily foraging behavior, followed protocols of Miller & Wagner (2014), which were based on standardized methods (Remsen and Robinson 1990). Between July 2009 and May 2014, we spent 295 field days throughout continental Australia, Kangaroo Island, and Tasmania (Fig. 1). When not driving between sites, we spent the daylight hours walking transects recording foraging movements, substrates, and food items. For each observation, we recorded the time, location, substrate on which the bird foraged, the attack maneuver employed, whether the bird was hanging during the maneuver, the height of the foraging bird, the height of the surrounding canopy, the distance of the bird from the trunk, and the density of foliage around the foraging bird. These last two variables were recorded on an ordinal scale. See the supplementary material for a complete list of all possible substrates, maneuvers, and details on foliage density and distances from the trunk.

**Figure 1.**
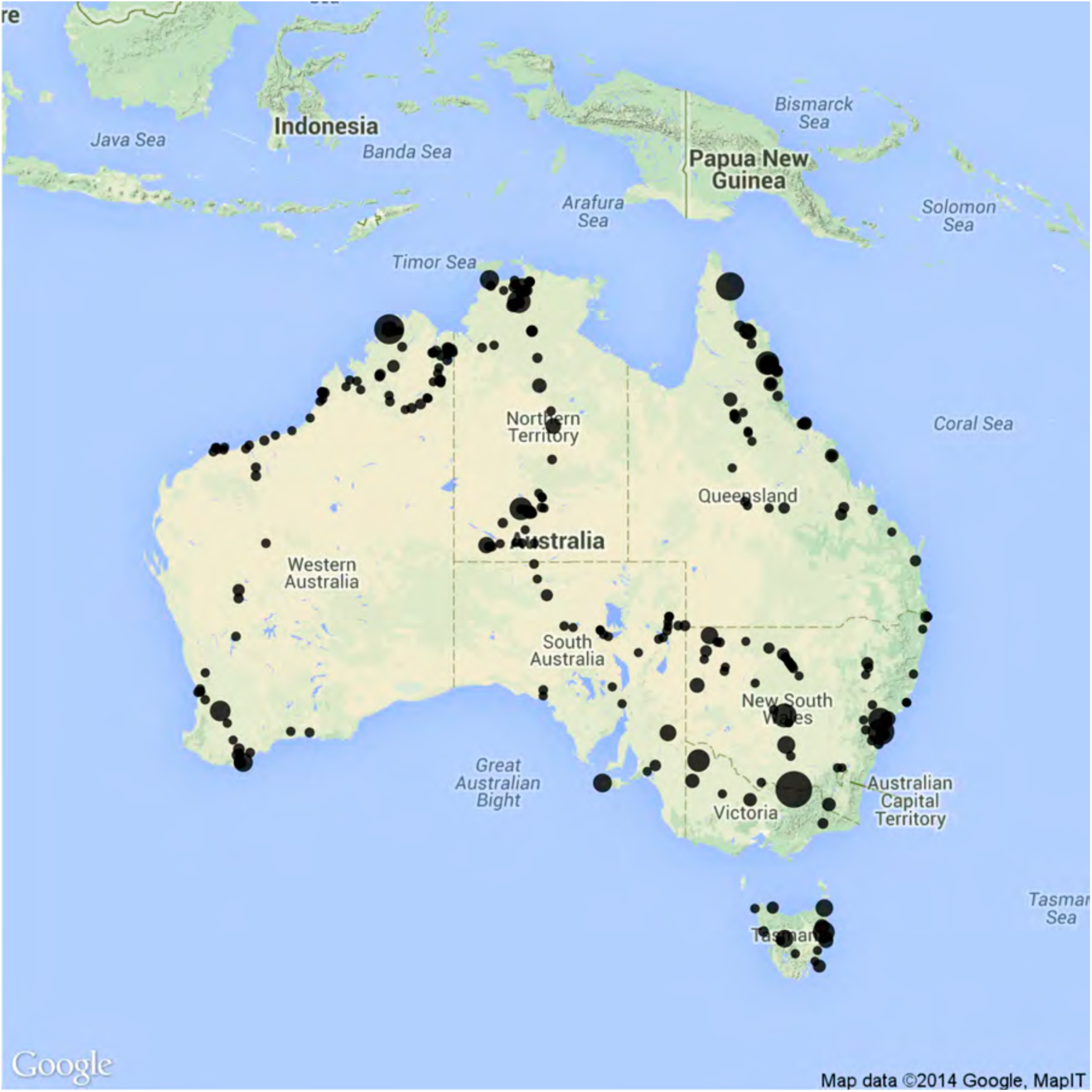
Map of Australia, showing the locations where foraging observations for this study were made. The size of the dot corresponds to the number of independent observations that were recorded at that site.

If the first foraging maneuver we observed initially drew our attention to an individual bird, we discarded the observation to minimize bias. Otherwise, if we located a bird, for instance, by its vocalizations, we recorded the first maneuver we saw. We endeavored to record only one observation per individual per day. To better understand individual variation in foraging behavior, in some cases we did record multiple observations from single birds. However, we considered such series of observations as collectively representing a single data point. We chose 20 independent observations as the minimum required for analysis of a species’ niche. We did not meet this requirement for the elusive Gray Honeyeater (*Conopophila whitei*), and therefore excluded it from ecological analyses.

### Meliphagidae assemblage and climatic data assembly

We used the species distribution dataset from Miller et al. (2013). This taxonomically and spatially cleaned dataset contains 2,273,404 localized observations of individuals across all Meliphagidae species. The data were downloaded and concatenated from eBird (Sullivan et al. 2009) and the Atlas of Living Australia (http://www.ala.org.au/). We defined assemblages as the species occurring within 100×100-km grid cells. In many parts of Australia it is a reasonable assumption that all species in a grid cell could interact ecologically. For instance, while collecting foraging data, we occasionally recorded a species list for the area. On average, we observed a mean of 40% of the bird species recorded from a given grid cell each day (*n* = 27, *SD* = 16%, max = 100%, min = 21%). We ran analyses using both the presence-absence community data matrix (CDM) and the relative abundance CDM which, to the extent that our assessment of abundance can be relied upon, can reduce the influence of vagrants and provide added biological detail on the effects of habitat filtering and competitive exclusion.

We used the mean annual temperature (MAT) and precipitation (MAP) layers from WorldClim (http://www.worldclim.org/bioclim), as summarized in Miller et al. (2013), to quantify variation in climate across the continent.

### Definition and summary of multivariate trait spaces

We log-transformed all linear morphological measurements (but not the composite indices), and then calculated species’ averages for all morphological and ecological measures. For ecological measures, species’ traits refer to mean average foraging height, proportion of attacks that were gleans, proportion of attacks that were to flowers, etc.

We used the R (R Development Core Team 2011) package *phytools* (Revell 2012) to ordinate each dataset separately with a phylogenetic correlation matrix-principal components analysis (pPCA, Revell 2009), i.e. a PCA where the Brownian-motion expected degree of covariance among species’ traits is incorporated into the calculation of PC axes and scores. To visualize how the Meliphagidae explored these trait spaces, we used a color-coded phylomorphospace approach (Miller et al. 2013), based on an updated, time-calibrated version of the Meliphagidae phylogeny for all analyses (Joseph et al. 2014), with 9 species added manually as in Miller et al. (2013) and Mast et al. (2015).

To quantify the phylogenetic signal in species’ morphological and ecological traits, we used the R package *geomorph* (Adams 2014). This approach incorporates the multivariate nature of these data, and instead of calculating a separate value per trait or principal component axis, outputs a single K value (Blomberg et al. 2003) that integrates all of the species’ traits simultaneously.

### Concordance of morphology and ecology

To examine the degree to which morphology predicts species’ ecologies, and vice versa, we used a phylogenetic canonical correlation analysis (pCCA, Revell and Harrison 2008). We interpreted the initial *phytools* results with custom scripts that calculate the phylogenetic generalized least squares (PGLS) correlation coefficients of the raw trait variables with species’ positions along the derived ecological and morphological canonical axes. We then used these correlation coefficients to calculate redundancy indices (Stewart and Love 1968) with the *candisc* package. These indices provide a measure of the amount of variance in the ecological dataset that can be explained with the morphological dataset, and vice versa. Many of the ecological variables are zero-skewed, reflecting the paucity of certain foraging behaviors, thus we repeated these analyses excluding the most zero-skewed behaviors. Because results were qualitatively identical with either ecological dataset, we do not discuss results from the reduced ecological dataset in detail.

### Calculation of assemblage trait disparity

We quantified (separately) the variation in morphological and ecological traits of species within each assemblage. To do this, we used species’ positions in multivariate trait space to calculate trait disparity as the mean pairwise distance (MPD_trait_) in Euclidean space among the points from a given assemblage, as defined in the community data matrices (CDMs). We did this both with the presence-absence and relative abundance CDMs (interspecific abundance-weighted MPD, Miller et al. 2015). A large value for this index signifies an assemblage composed of species that are, on average, widely separated from each other in trait space, while a small value signifies an assemblage of species that are clustered in trait space. When used with relative abundance data, common species contribute more to MPD_trait_, and distances between such species more strongly influence the metric.

To account for the potential influence of dispersal limitation on community structure, we calculated standardized effect sizes (SES) of these trait distances, after 1,000 randomizations, using a new dispersal null model. This model constructs CDMs where assemblage species richness, individual species’ occurrence frequencies, and total CDM abundance (i.e., total number of individuals in the CDM) are maintained, and it settles species in these assemblages with a probability proportional to their relative abundance in nearby cells, thereby incorporating dispersal probabilities directly into expectations. We used the product of the geographic distance matrix (great circle distances) and the environmental distance matrix (Euclidean distance after correlation matrix PCA) as our measure of distance among grid cells. These distances are standardized scores that reflect observed assemblage deviation from expected trait packing metrics (MPD_trait_) given realistic assembly processes. As an additional check on these results, in the supplementary material we also derived SES using a null model that maintains observed species richness but not occurrence frequencies or dispersal probabilities.

To compare how these standardized trait disparity scores scale with phylogenetic diversity, we calculated similar SES for mean pairwise phylogenetic distance (MPD_phylo_) using both the new null model and, in supplementary material, the richness only model. Like the SES for trait disparity, large SES values here reflect assemblages that are composed of species that are evenly spread across the phylogeny, compared to null expectations.

### Species-based measures of trait evolution

To calculate species-specific rates of trait evolution, we built on functions from *convevol* (Stayton 2015) to calculate the distance evolved, along the inferred evolutionary branches, of each species through multivariate trait space from the inferred root. The resulting “traits” represent the total distance evolved by a given species (and its ancestors) from the most recent common ancestor (crown node) of the honeyeaters. Thus, large values reflect species that are deduced to have undergone notable phenotypic shifts over their evolutionary history. We then input these species-specific “traits” into a trait-diversification analysis in BAMM (Rabosky 2014; Rabosky et al. 2014), and took the tip-averaged trait diversification rates as our measure of rate of trait evolution away from the root of the Meliphagidae. This method tests whether certain species, such as those from the desert, have evolved through trait space at a faster rate than others.

We developed a method to quantify how different a species is, phylogenetically or in trait space, from the numerous species it occurs with across its range. The goal was to provide a species-centered examination of the prediction that species from phylogenetically clustered assemblages have evolved through trait space at a faster rate than those from less phylogenetically clustered assemblages. Building upon the work of Villalobos et al. (2013), we extended the phylogenetic field concept to include standardized effect sizes (SES) and abundance-weighting. Thus, we defined a species’ phylogenetic field as the weighted mean of either the non-abundance-weighted (hereafter, “unweighted”) or interspecific abundance-weighted MPD_phylo_ of the grid cells it occurred in (Miller et al. 2015). The weights used in this species-specific measure were that species’ relative abundance in those grid cells (when using unweighted MPD, this reduces to a standard, unweighted mean). Similarly, we defined a species’ trait field as the weighted mean of the ecological or morphological trait neighborhood (MPD_trait_) it occurs in across its range. We defined the standardized form of these trait and phylogenetic fields by deriving 1,000 CDMs with the dispersal null model (and 1,000 CDMs with the richness null in supplementary material), and then used these to calculate simulated species’ fields. We used these values to calculate field SES, per species, as the difference between the observed field and the mean of the simulated fields divided by the standard deviation. Thus, for each species, this metric measures the average properties of local assemblages in which that species occurs. For instance, species with large trait field SES values occur in assemblages with species that are evenly distributed in trait space, after accounting for dispersal probabilities and observed species richness. Scripts to calculate phylogenetic and trait fields, including the standardized versions of these, have been included in the *metric Tester* package (Miller et al. 2015).

### Correlating patterns of trait evolution, trait community structure, and climate

To test our prediction that Meliphagidae trait disparity does not parallel phylogenetic diversity, we used ordinary least squares (OLS) regression to compare the standardized morphological and ecological trait disparity scores from each grid cell with their underlying MAT and MAP.

To further examine any disconnect between trait and phylogenetic diversity, we explored rates of trait evolution and their potential drivers. Taking a species-specific perspective, we used PGLS regressions to test whether rates of morphological or ecological trait evolution from the BAMM analysis were correlated with a species’ phylogenetic field. This approach considers species individually, and does not consider whether the co-occurring species are evenly partitioning trait space (e.g., it is possible that all species in an assemblage could have evolved quickly towards the same trait combination). Thus, we also used PGLS to compare species’ trait fields with their phylogenetic fields. This approach more directly compares how a species partitions niche space among its potential competitors as a function of the phylogenetic neighborhood it finds itself in. For example, if species in phylogenetically clustered assemblages were widely separated in trait space, then, despite close phylogenetic relationship, such species would be evenly partitioning niche space with their potential competitors.

## RESULTS

### Summary of the datasets

We measured 710 specimens of 74 of 75 Australian Meliphagidae species, although we did not take the complete set of measurements on each specimen. Sample sizes range from 1 for the range-restricted White-lined Honeyeater (*Meliphaga albilineata*), to 36 for White-plumed Honeyeater (*Ptilotula penicillata*). The species-averaged dataset will be available upon publication of the final version of this manuscript.

We collected 9,595 foraging observations across 74 species of Australian honeyeater. After accounting for serial observations, the dataset contains 7,302 independent observations. The most-observed species was Brown Honeyeater (*Lichmera indistincta*, n=459). The least-observed species was Green-backed Honeyeater (*Glycichaera fallax*, n=20). The one individual observed of Gray Honeyeater (*Conopophila whitei*) was excluded from analysis. The species-averaged dataset provides detailed, quantitative measures of the foraging ecology of a large continental radiation of vertebrates, and will be available upon publication of the final version of this manuscript.

### Multivariate trait spaces

The first three axes (out of 15) from the morphological PCA captured 78% of the variance in the dataset. The first described differences in overall body size. The second described an axis of variation from species with long, decurved bills to those with nares positioned towards the middle of the bill, and bills whose depth and width taper considerably over their length. The third axis separated species with rounded wings and long bills that taper in depth (e.g., Eastern Spinebill, *Acanthorhynchus tenuirostris* and Tawny-breasted Honeyeater, *Xanthotis flaviventer*) from those with pointed wings and bills with much of the length proximal to the nares (e.g., Regent Honeyeater, *Anthochaera phyrgia* and Yellow-throated Honeyeater, *Nesoptilotis flavicollis*). Multivariate K for the morphological dataset was 0.863 (*p* = 0.001 that K differs from 0), emphasizing that species show a strong tendency to resemble their ancestors (Fig. 2).

**Figure 2.**
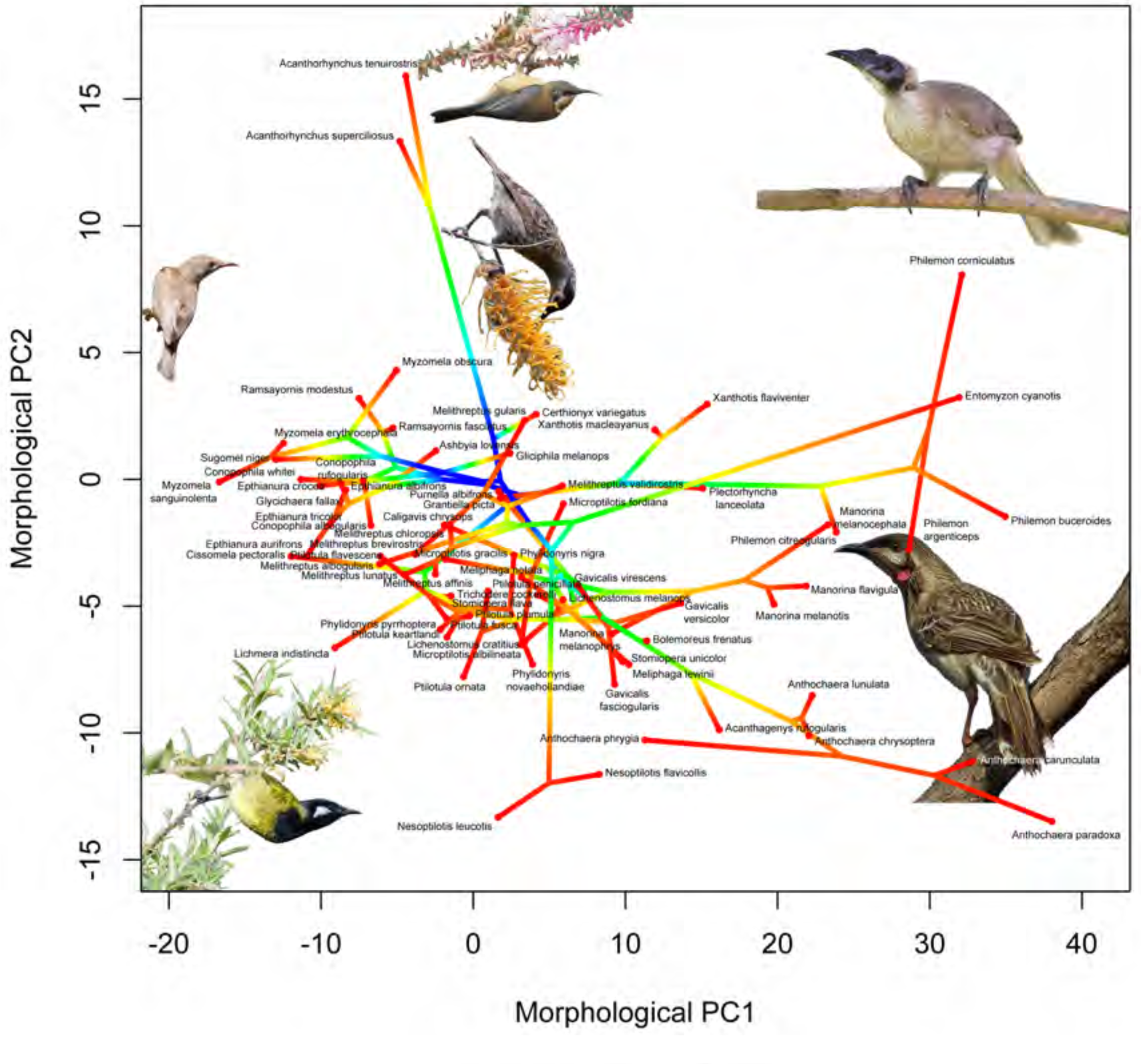
Phylomorphospace showing the first two axes from the phylogenetic, correlation matrix-principal components analysis on the morphological data. Time since the root (~20 mya) is colored from blue to red. The first axis represents a general size axis, with larger species on the right. From top to bottom, the second axis separates species with decurved bills from those with bills that taper considerably over their length in depth and width. These first two axes account for 68% of the variance in the dataset. In clockwise order from the top right corner, photographs are of Noisy Friarbird (*Philemon corniculatus*), Red Wattlebird (*Anthochaera carunculata*), Whie-eared Honeyeater (*Nesoptilotis leucotis*), Brown-backed Honeyeater (*Ramsayornis modestus*), Eastern Spinebill (*Acanthorhynchus tenuirostris*), and Macleay’s Honeyeater (*Xanthotis macleayanus*). The *Ramsayornis* and *Xanthotis* photos are by Bryan Suson; all others used under a Creative Commons license.

Of the 30 axes from the ecological pPCA, the first three described 37% of the variance in the dataset (the first 10 described 76%). With the reduced dataset (17 variables) the first three axes described 54% of the variance. Returning to the full 30-axis phylogenetic pPCA, the first principal component described an axis of variation from highly nectarivorous species that, when not foraging on flowers, tended to sally-strike for flying invertebrates, to species that gleaned frequently from leaves and branches. The second axis distinguished species that foraged among foliage and took food from amidst leaves, from species that foraged more in the open and took food from hanging bark or branches, and employed a rare foraging maneuver called gaping, whereby the bill is inserted into a substrate such as a rolled leaf or under bark and levered up to pry open the substrate. The third principal component differentiated species that foraged high in the canopy and those that foraged on the ground, well away from the cover of trees. Multivariate K for the ecological dataset was 0.471 (*p* = 0.002), emphasizing that while many species resemble their relatives in foraging behavior, others have become differentiated by evolution across considerable ecological distances (Fig. 3).

**Figure 3.**
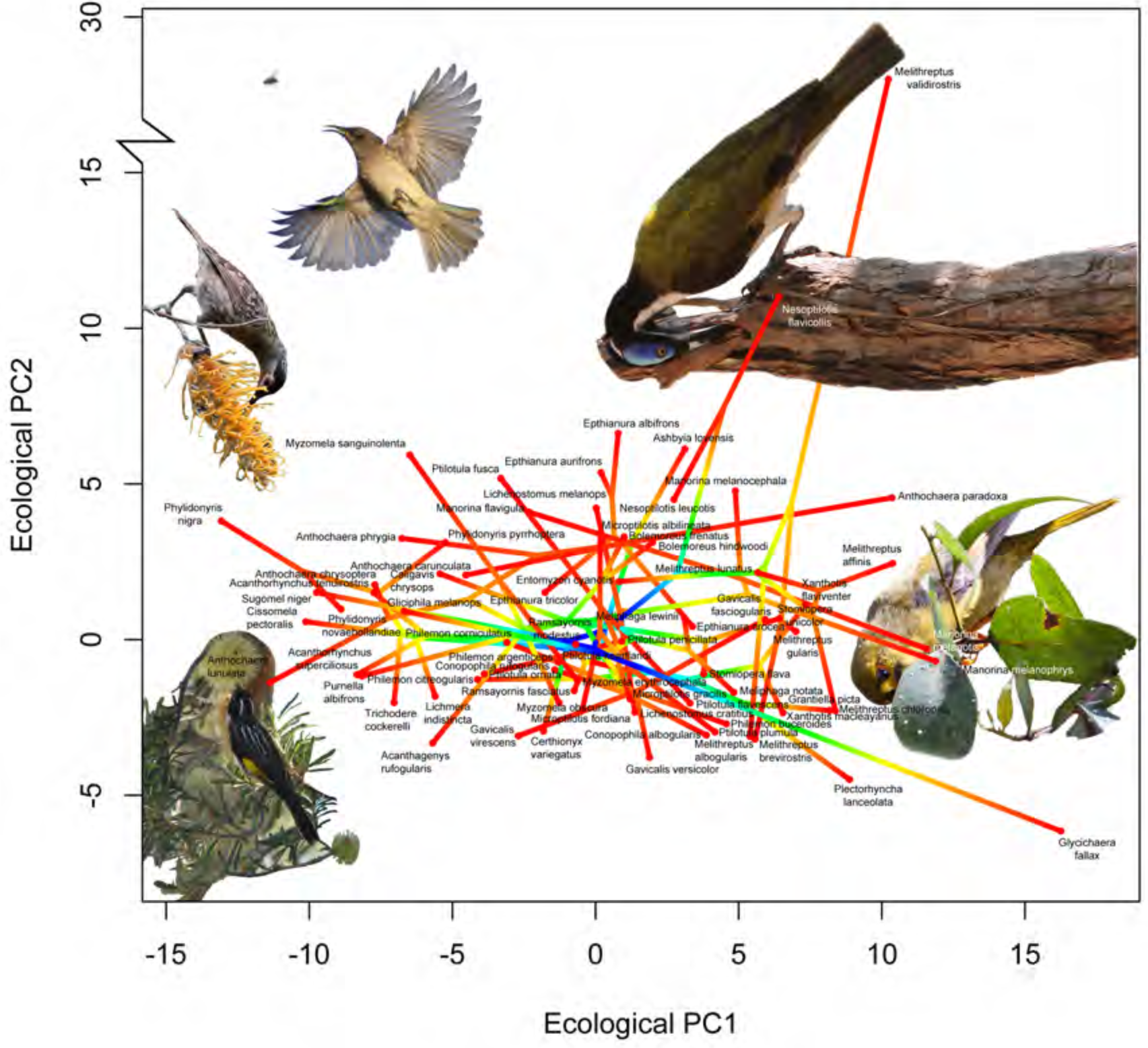
Phylomorphospace showing the first two axes from the phylogenetic, correlation matrix-principal components analysis on the ecological data. Time since the root (~20 mya) is colored from blue to red. From left to right, the first axis separates highly nectarivorous species that, when not foraging on flowers, tend to sally-strike for aerial invertebrates, to species that glean frequently from leaves and branches. From top to bottom, the second separates species that forage more in the open and take prey from hanging bark, branches, and employ a rare foraging maneuver called gaping, to those that forage in leafy situations and take prey from leaves. Not shown is the extreme position occupied by species of *Epthianura* and *Ashbyia* on the third axis. These first two axes account for 28% of the variance in the morphological dataset. Note the broken y-axis/extreme position occupied by Strong-billed Honeyeater (*Melithreptus validirostris*). Photographs are chosen to illustrate behaviors, rather than the species themselves. In clockwise order from the top right corner: Blue-faced Honeyeater (*Entomyzon cyanotis*) probing a branch, Bell Miner (*Manorina melanophrys*) gleaning lerp from a leaf, New Holland Honeyeater (*Phylidonyris novaehollandiae*) probing a *Banksia marginata* flower in dense vegetation, Macleay’s Honeyeater (*Xanthotis macleayanus*) probing an exposed *Grevillea pteridifolia* flower, and Brown Honeyeater (*Lichmera indistincta*) sally-striking for an insect. The *Entomyzon* and *Xanthotis* photos are by Bryan Suson; all others used under a Creative Commons license.

### Canonical correlation analysis

The first four axes of the pCCA were statistically significant (Table 1). Collectively, the morphological dataset explained 37% of the variance in the ecological dataset. The ecological dataset explained 62% of the variance in the morphological dataset. Proportions of variance explained were 39% and 41% with the reduced ecological dataset. The first canonical variate described an axis ranging from species with decurved bills and pointed wings that are highly nectarivorous and regularly sally-strike for aerial invertebrates to those with long tarsi and toes, wide and deep bills, and heavy mass that glean, forage on the ground and on branches, and employ pecking and sally-pouncing (Table 2). The second described an axis from species with long bills and tarsi that frequent flowers with long corollas and often take insects in aerial pursuits, to those that have short tarsi, pointed wings, and hang while foraging in tall canopies. The third described a tradeoff between species with bills that taper considerably in depth and forage on branches to those with deep, decurved bills that forage high in the canopy, often from hanging bark, and glean fruits. The fourth described an axis from species with bills that taper considerably in depth, employ pecking maneuvers, and forage on the ground, well away from trees, to species with long tails, overall long body length, pointed wings and a considerable portion of their bill proximal to the nares, and forage relatively high in the available canopy, often on branches and on flowers with long corollas, and often hang to do so (Table 2).

### Assemblage trait disparity and species richness and climate correlates

Morphologically, most Meliphagidae assemblages did not deviate beyond statistical expectations given the dispersal null model, which simulated realistic assembly processes (i.e. their SES were between −1.96 and 1.96). The same was also true given a null model that only maintained observed species richness (supplementary material). With presence-absence data, none of 695 assemblages were significantly clustered in trait space, while only one was significantly overdispersed. When calculations were species abundance-weighted, one site was significantly clustered and nine were overdispersed. Morphological SES for MPD_trait_ increased slightly along a gradient of increasing species richness whether abundance-weighted (*r*^2^ = 0.097, *p* < 0.001) or not (*r*^2^ = 0.057, *p* < 0.001). Thus, species rich sites were composed of morphologically more evenly spaced species.

Morphological SES MPD_trait_ decreased slightly along a gradient of increasing temperature (unweighted *r^2^* = 0.012, *p* < 0.001; abundance-weighted *r^2^* = 0.036, *p* < 0.001), and increased slightly along a gradient of increasing precipitation (log_10_(MAP), unweighted *r^2^* = 0.154, *p* < 0.001; abundance-weighted *r^2^* = 0.296, *p* < 0.001). Thus, the coldest, wettest sites contained species that were most evenly spread in morphological trait space, although this pattern was weak.

As in the case of morphological diversity, most Meliphagidae assemblages were not significantly structured in ecological space with the dispersal null model. When unweighted, six (of 695) assemblages were clustered and none were significantly overdispersed. When abundance-weighted, 25 were clustered and none were overdispersed. Approximately 10% of sites were considered significantly clustered with the simple richness null model (supplementary material). Unlike morphological trait disparity, when unweighted, ecological trait disparity was unrelated to species richness (*r*^2^ < 0.001, *p* = 0.501), and when abundance-weighted, ecological SES MPD_trait_ was negatively correlated with species richness (*r*^2^ = 0.196, *p* < 0.001). Thus, the most abundant species in the most species-rich sites were often ecologically similar.

Ecological trait disparity decreased slightly along gradients of increasing temperature when unweighted (*r*^2^ = 0.066, *p* < 0.001, relationship not significant when abundance-weighted). Ecological SES MPD_trait_ was also negatively correlated with precipitation (unweighted *r^2^* = 0.127*, p* < 0.001; abundance-weighted *r^2^* = 0.013*, p* = 0.003). Thus, the coldest, driest sites tended to contain species that were most evenly spread in ecological trait space, although we emphasize that the overriding signal here was the absence of a strong pattern.

In short, Meliphagidae trait disparity does not closely parallel phylogenetic diversity (Fig. 4). Only weakly significant positive correlations exist between morphological trait disparity and phylogenetic diversity (unweighted *r^2^* = 0.174, *p* < 0.001; abundance-weighted *r^2^* = 0.291, *p* < 0.001), and between ecological trait disparity and phylogenetic diversity (unweighted *r^2^* = 0.015, *p* = 0.001; abundance-weighted *r^2^* = 0.112*, p* < 0.001).

**Figure 4.**
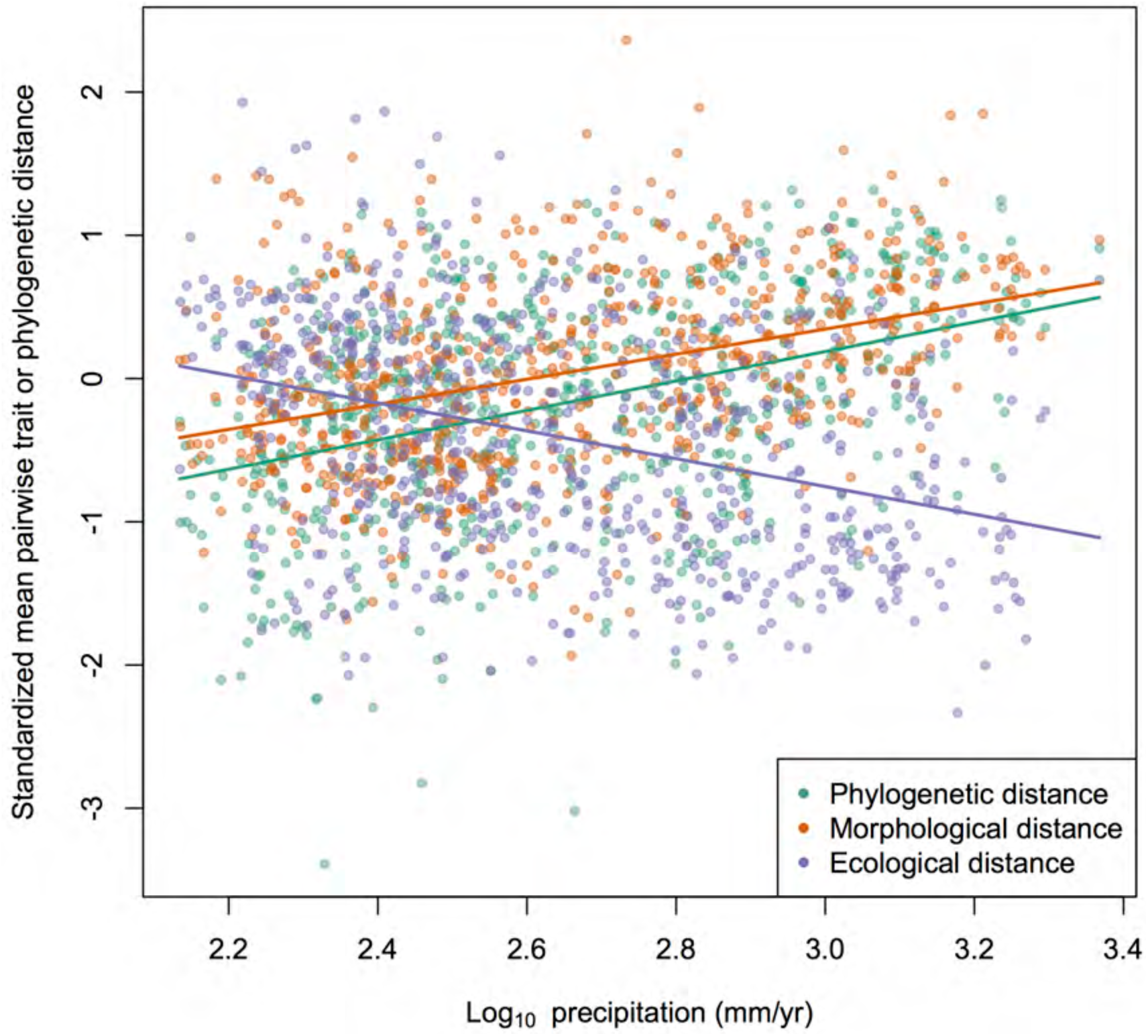
Standardized phylogenetic diversity and morphological and ecological trait disparity, per assemblage, as compared with the logged mean annual precipitation at the site. These mean pairwise phylogenetic distances and mean pairwise Euclidean distances in multivariate trait space were standardized with a null model that reflects dispersal probabilities and maintains species richness and occurrence frequencies in the simulated assemblages. While morphological disparity closely mirrors phylogenetic diversity, declining towards the arid interior, ecological disparity actually increases along the same gradient. Species in the desert, while closely related and similar looking, actually forage in dramatically different ways given their close affinities.

### Rates of trait evolution

Considering distance evolved through trait space, the estimated sample size for all BAMM runs exceeded the recommended minimum of 200 (minimum used = 495) for both the potential number of shifts and the log-likelihood.

For morphological traits, the single best shift configuration included an increase in the rate of evolution of the wattlebird clade (*Anthochaera*, excluding *Acanthagenys*), and an otherwise overall declining rate of trait evolution (occurred in 23% of runs). Seven percent of the runs localized the *Anthochaera* shift one node deeper in time, in the clade that also includes Spiny-cheeked Honeyeater (*Acanthagenys rufogularis*), and 6% placed it one node deeper than that, in the clade that also includes Bridled Honeyeater (*Bolemoreus frenatus*; B. *hindwoodi* was not sampled). Another configuration, observed in 8% of runs, included three increases in rates of morphological trait evolution: in the *Anthochaera* clade, on the branch leading to Blue-faced Honeyeater (*Entomyzon cyanotis*), and in the friarbird clade (*Philemon*). All other likely shift configurations were variations on this general theme; that is, they included increases in the rates of evolution of branches leading towards larger-bodied species.

For ecological traits, the single best shift configuration included no abrupt shifts in the rate of trait evolution, and a phylogeny-wide rate of trait evolution that declined continuously through time (frequency of occurrence 58%), suggesting that ecological space was filled relatively early in the evolution of the group. However, 18% of runs did include an increase in the rate of evolution of the Strong-billed (*Melithreptus validirostris*) + Brown-headed (*M. brevirostris*) + Black-chinned Honeyeater (*M. gularis*) clade (= *Eidopsarus*, Toon et al. 2010), 14% localized that shift on the Strong-billed branch, 5% placed the increase in rate of evolution one node deeper, on the clade that also included White-throated Honeyeater (*M*. *albogularis*), and 2% placed it at the base of the entire genus. These seven *Melithreptus* species, particularly those of the *Eidopsarus* subgenus, frequently probe branches and are notably less nectarivorous than most other Meliphagidae.

### Phylogenetic and trait fields

After accounting for dispersal probability based on environmental and geographic distances, species tended to occur in overdispersed phylogenetic fields. When unweighted, mean SES MPD_phylo_ was 5.45 (Wilcoxon signed rank test *p* < 0.001). When abundance-weighted, the mean was 3.13 (*p* = 0.001). With the richness null model, which does not account for dispersal probability, species tended to occur in clustered phylogenetic fields (supplementary results). Morphologically, species were variable in terms of the trait fields they occurred in, ranging from SES = −10.89 to 10.22 when unweighted (mean = 0.01, Wilcoxon test that *μ* = 0, *p* = 0.43). Species were similarly variable when MPD_trait_ was abundance-weighted, ranging from −15.75 to 18.35, but there was a significant trend towards overdispersion (mean 3.45, *p* < 0.001).

From an ecological perspective, when unweighted, species tended to occur in overdispersed trait fields, with SES values that ranged from −5.93 to 23.98 (mean 5.79, *p* < 0.001). However, when abundance was incorporated, most species occurred in ecologically clustered assemblages, with SES values that ranged from −13.95 to 15.08 (mean −2.17, *p* < 0.001). In other words, considering only presence and absence across their range, focal species tended to occur with ecologically disparate species. When also incorporating abundance, focal species tended to occur with abundant and ecologically similar species (presumably because there is a trend for ecologically outlying species to be less common than species towards the center of niche space). Additional trait field results, including both unstandardized values and those standardized with the richness null model, are presented in supplementary material.

### Potential drivers of rates of trait evolution and variation in trait fields

As noted above, little variation was found in species’ rates of trait evolution; these rates were not related to species’ phylogenetic fields.

Species’ standardized morphological trait fields, however, were closely correlated with their phylogenetic fields. Species from less phylogenetically clustered assemblages tended to occur in assemblages that were less clustered in morphological trait space (unweighted PGLS *r^2^* = 0.20, *p* < 0.001; abundance weighted *r^2^* = 0.49, *p* < 0.001). In contrast, the relationship between species’ standardized phylogenetic and ecological trait fields was only weakly positive (unweighted PGLS *r^2^* = 0.001, *p* = 0.84; abundance weighted *r^2^* = 0.07, *p* = 0.02). When species’ raw (i.e., unstandardized), unweighted ecological trait fields were compared to their raw, unweighted phylogenetic fields, species from the most phylogenetically clustered assemblages tended to occur in assemblages that were the least clustered in ecological trait space (supplementary material); these species not only tended to be dissimilar from the species they co-occur with, but they tended to occupy positions on the periphery of ecological space.

## DISCUSSION

The Meliphagidae, or honeyeaters, are a diverse family of passerines distributed predominantly in Australia, New Guinea, and the Pacific Islands. They occupy a wide range of ecological regions, with at least one species occurring almost everywhere in Australia, including Tasmania, where there are four endemic species. Most species take some nectar, but some are highly frugivorous, and others are dedicated insectivores (Higgins et al. 2001). Owing both to ease of observation, and a history of interest in these species in Australia, their foraging behavior has been studied in some detail (Recher 1971; Ford and Paton 1976a; Paton 1980; Pyke 1980; Ford and Paton 1982; Recher et al. 1985; Ford 1990). These studies laid the foundation upon which this paper is based.

Given the diversity of resource acquisition strategies in the honeyeaters, whether well-defined ecomorphological relationships would emerge from our analysis was unclear. For instance, the external morphological characters that distinguish small-fruit-eating passerines from insectivores are not always evident. Moreover, owing to patchy resources, unpredictable flowering phenology and many short-corolla, generalist-accessible flowers, the Australian honeyeaters are considered uniquely unspecialized in their floral preferences when compared to groups like the hummingbirds (Paton and Ford 1977; Stiles 1981). Despite this, our expectation that morphology predicts ecology in the group was borne out by the phylogenetic canonical correlation analysis (pCCA). Results from the pCCA provide numerous insights into ecomorphological relationships within the honeyeaters, and we briefly discuss a few of these below.

As noted above, honeyeaters are considered unspecialized in their floral preferences (Paton and Ford 1977). After many field hours, however, we knew this to be an overstatement, with certain species like Eastern Spinebill (*Acanthorhynchus tenuirostris*) showing marked bill-flower matching (Fig. 5; indeed, Paton and Ford allude to potentially higher specificity in this and a few other species). Results from the pCCA clearly show a dataset-wide correspondence between the length of species’ bills and the flowers they visited. Matching between honeyeaters and particular floral resources extends beyond the “specialized” species like spinebills (pers. obs.), and additional investigation into honeyeater-plant networks is warranted. Nevertheless, the overall degree of matching does appear decidedly lower than that in hummingbirds. Nectar is generally abundant in Australia (Orians and Milewski 2007). For instance, some clearly ornithophilous flowers (e.g., *Grevillea speciosa*, Fig. 5) have distinct slits in the base of the floral tube, rendering nectar readily available to even such short-billed species as Brown-headed Honeyeater (*Melithreptus brevirostris*, pers. obs.). Indeed, the most important nectar resources across Australia are probably the cup-like flowers of *Eucalyptus* (Woinarski et al. 2000).

**Figure 5.**
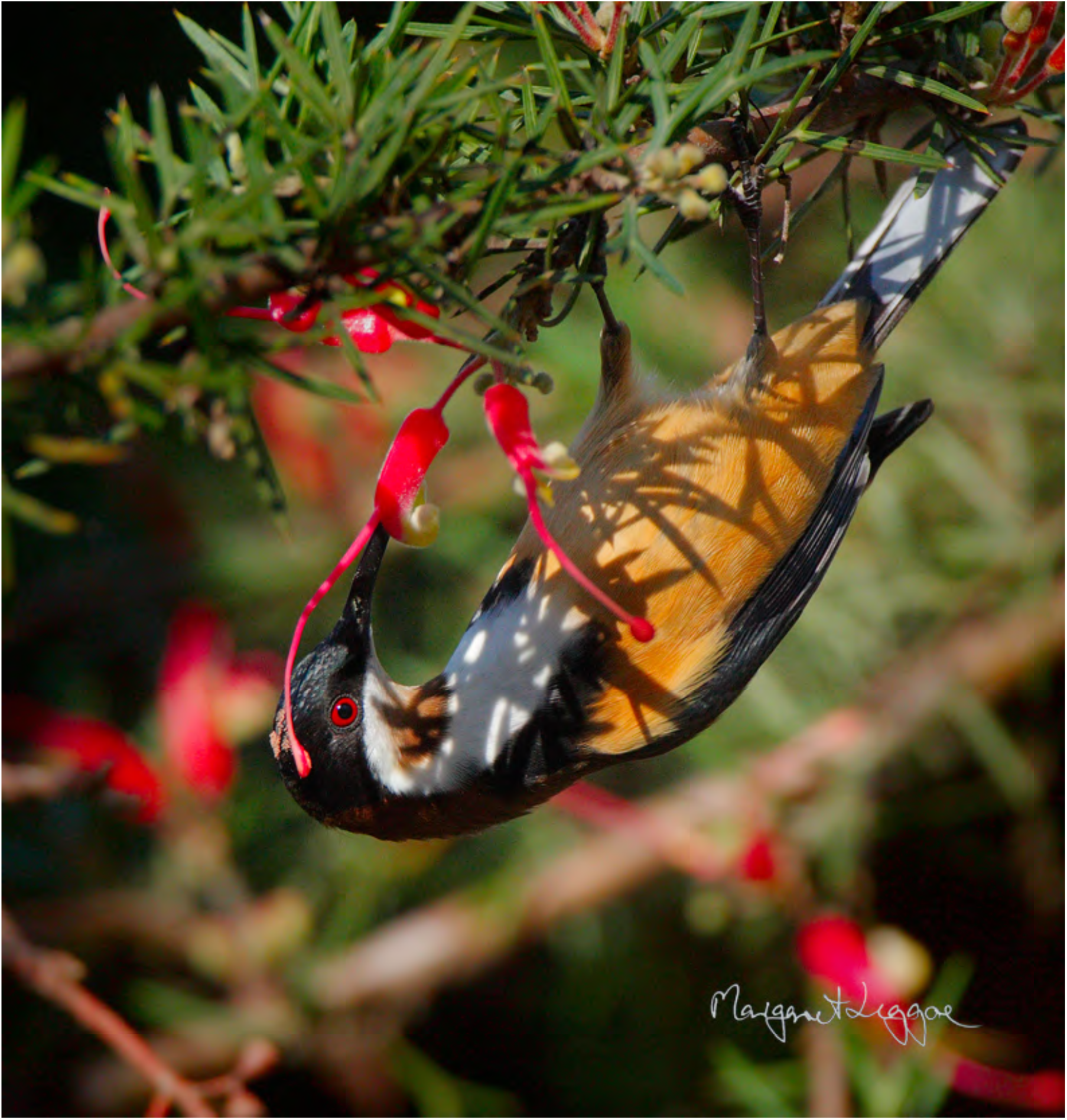
An Eastern Spinebill (*Acanthorhynchus tenuirostris*) hangs upside-down to probe a *Grevillea tripartita tripartita*. Pollen daubed on the spinebill’s forehead by the pollen presenter is clearly visible. Many *Grevillea* species have long tubular corollas and a clear morphological matching between long-billed honeyeater species like the spinebill yet, owing to a slit in the floral tube, many species are also readily accessed by short-billed species like Brown-headed Honeyeater (*Melithreptus brevirostris*). Photo by Margaret Leggoe.

Among hummingbirds, long bills are associated with feeding on flowers with long corollas (Snow and Snow 1980), while among flycatchers long bills are associated with aerial attacks on invertebrates (Fitzpatrick et al. 2004). Though this may seem like phenotypic convergence towards dramatically different ecologies, in the honeyeaters, we found a close relationship between degree of nectarivory, aerial attacks and bill length. That some of the most nectarivorous honeyeaters regularly “hawk” (make aerial attacks for flying invertebrates) is well known (Recher and Abbott 1970). Preliminary analyses suggested such species have a surfeit of calories in the form of nectar, and that these energetically costly maneuvers are used to rapidly procure protein to supplement their otherwise protein-poor primary food resource (Ford and Paton 1976b). These studies, however, were limited to four species. The continental, cross-season, 74-species, dataset-wide trend found here suggests the existence of a more fundamental dimension of variation (Westoby et al. 2002), from species that move steadily through foliage, gleaning invertebrates and occasionally taking some nectar, to those that take much nectar, and only occasionally perform energetically costly hawking maneuvers in pursuit of protein. Hummingbirds are also well known to occasionally supplement their diet with flying invertebrates; indeed, their beaks actually deform to facilitate aerial insect capture (Yanega and Rubega 2004).

The species on the periphery of ecological trait space, those that drove patterns of assemblage structure, tended to be well-known ecological eccentrics. However, at least according to our measurements, these species were not notably deviant in morphospace, and it is with these species that the recovered ecomorphological relationships begin to break down. Thus, Painted Honeyeater (*Grantiella picta*), a species that foraged on fruit 47% of the time in our data (almost twice as frequently as the next most frugivorous species), resembles Gilbert’s Honeyeater (*Melithreptus chloropsis*), a species that was never observed to eat fruit. And chats of *Epthianura* and *Ashbyia*, long considered a separate family due to their idiosyncratic, open-country foraging behavior, differ little in gross morphology from *Conopophila*, *Ramsayornis*, and *Glycichaera*. Finally, the genus *Melithreptus*, including the single largest ecological outlier, Strong-billed Honeyeater (M. *validirostris*), overlaps in morphospace with other genera. Our morphological measures did not capture any aspects of internal morphology (Ricklefs 1996); chats, for instance, have lost most of the bristles on their brush-tipped tongues (Parker 1973). However, even if we had measured such traits, it is likely that some of the recovered ecomorphological relationships are particular to the Australian honeyeaters. Whether such relationships will translate to other avian groups remains to be tested; the family Nectariniidae would be of particular interest in this regard.

Another internal morphological character with clear ecological ramifications is a unique jaw articulation in some species, particularly Strong-billed Honeyeater, other *Melithreptus*, and Black-eared and Yellow-throated Miners (*Manorina melanotis* and *M. flavigula*) (Bock and Morioka 1971). This articulation, concealed in traditionally prepared specimens, was postulated to facilitate raising the maxilla when force was applied to it, allowing the tongue to be moistened with saliva and then extruded from the bill. However, these authors were unsure of the precise use for such morphology, and while they cited Keast (1968), they apparently did not notice his report therein of what we call gaping, i.e., using the bill to pry apart vegetation (Remsen and Robinson 1990) in Strong-billed Honeyeater. Others have discussed the potential use of this articulation in *Melithreptus* (Willoughby 2005), but in general its behavioral repercussions remain poorly studied. In our dataset, the correlation between the articulation and gaping was quite clear. The species that most frequently employed gaping was Strong-billed Honeyeater. Three other species of *Melithreptus* also used the technique, as did Black-eared and Yellow-throated Miners. However, the sister to *Melithreptus*, Blue-faced Honeyeater (*Entomyzon cyanotis*) also employed gaping occasionally, as did both species of *Stomiopera*, particularly White-gaped Honeyeater (*S*. *unicolor*). Bock and Morioka (1971) examined jaws of Blue-faced Honeyeater, finding them to be devoid of the articulation, and they also examined an unspecified number of species of *Meliphaga sensu lato* (to which *Stomiopera* used to belong), and similarly found no sign of the jaw gaping morphology. Whether they studied *Stomiopera* skeletons is unclear. It seems likely that careful study of *Stomiopera* will reveal similar jaw articulations to those seen in *Melithreptus* and *Manorina*.

Meliphagidae assemblages are not notably structured in ecological trait space, and ecological space filling does not vary across climatic gradients. This stands in stark contrast to how phylogenetic structure and morphological trait space of Meliphagidae assemblages varies along climatic gradients, where the arid interior is characterized by phylogenetically clustered (Miller et al. 2013) assemblages of morphologically similar honeyeaters. Thus, over evolutionary timescales, over large geographic and climatic spaces, and perhaps in conjunction with shorter time-scale competitive exclusion processes, honeyeaters have diversified to fill comparable local trait spaces in different climatic regions. This is particularly notable when considered in light of the close phylogenetic relationships among desert species.

How did honeyeaters accomplish this impressive filling of trait space? According to our BAMM analysis, it does not appear that species from phylogenetically clustered assemblages have systematically evolved any faster through ecological or morphological trait space. Of course, evolutionary rates are distinct from directions. Species in phylogenetically clustered assemblages might have exhibited directional evolution away from potential competitors. Our comparisons of species’ phylogenetic fields and their trait fields provide indirect evidence of this possibility. As one would expect, species’ phylogenetic and morphological trait fields were positively related. That is, species that occur in the most phylogenetically overdispersed assemblages also occur in the most morphologically overdispersed assemblages. Yet, this relationship was not manifested from an ecological perspective. Thus, after accounting for variation in species richness and for dispersal limitation, species from phylogenetically clustered assemblages are just as evenly arrayed ecologically as species from much more phylogenetically diverse assemblages. And, when unstandardized, the relationship between species’ ecological trait and phylogenetic fields was actually strongly negative (supplementary material). This suggests that despite close phylogenetic relationships and a tendency towards similar morphologies, the species from arid regions have diverged in ecology to a degree that they are, on average, as different from their competitors in foraging ecology as are those species that co-occur in mesic areas.

Our results are based on an ecological space that is defined by species’ foraging traits. A potentially more informative approach would define that ecological space *a priori*, with independent measures of local resource availability. By simulating community assembly where the settled species are drawn from ecologically similar sites, our dispersal null model goes part of the way towards addressing this shortcoming. Previous ecomorphological studies have included comparatively small numbers of species (e.g., Saunders and Barclay 1992), phylogenetically disparate/uneven comparisons (e.g., Douglas and Matthews 1992), single-site comparisons (e.g., Miles and Ricklefs 1984), and, owing to the difficulty of obtaining quantitative resource-use measures, fairly gross descriptors of species’ ecology (e.g., Pap et al. 2015). We avoided these issues by examining ecomorphological relationships in a phylogenetic context across a large clade over continental spatial scales. Our approach is admittedly confronted by spatio-temporal limitations, and we hope that citizen scientists may ultimately help to contribute sufficient data to address these questions over even larger scales.

Based on the results shown here, we conclude that species in the arid interior, those in phylogenetically clustered assemblages, have not evolved any faster through trait space. Instead, they have diverged from each other so as to partition trait space to an equivalent degree to that seen in more mesic areas. Morphology predicts ecology in the Australian Meliphagidae. However, the relationships show marked flexibility. Certain lineages, like the chats, forage in radically different ways from their relatives, exploiting entirely divergent resources with fairly conserved morphologies. Thus, community assembly and trait diversification in the honeyeaters reflects a combination of adaptation, both to local habitats and to competitors, and constraint as a result of past evolutionary history.

## ACKNOWLEDGEMENTS

ETM is grateful for financial support from the National Science Foundation, the St. Louis Audubon Society, a Trans-World Airlines Scholarship from the University of Missouri, and Macquarie University Higher Degree Research Office. We thank Leo Joseph, Árpád Nyári, Alicia Toon, Brian Venebles, Stephen Murphy, Hugh Ford, Harry Recher, Phillip Maher, Mick Roderick, Dick Cooper, David Watson, Keith and Lindsay Fisher, Alex Anderson, Bryan Suson, Josef Uyeda, Mark Westoby and Amy Zanne for invaluable discussion and advice on study sites, identification tips, and help in the field. We thank the FMNH, John Bates and Dave Willard, the AMNH, Paul Sweet, Santiago Claramunt and Joel Cracraft, the Burke Museum and Rob Faucett, and the Australian Museum, Walter Boles and Jaynia Sladek, for facilitating study at those institutions. We thank Margaret Leggoe for permission to use the photograph of the spinebill.

---

**Table 1.** Canonical correlations, chi-square values and significance of the correlations for the 15 dimensions from the phylogenetic canonical correlation analysis comparing Meliphagidae ecology and morphology.

| Dimension | Canonical correlation | Chi-square | p-value |
| --- | --- | --- | --- |
| 1 | 0.97 | 695.73 | < 0.001 |
| 2 | 0.93 | 549.28 | < 0.001 |
| 3 | 0.89 | 453.17 | < 0.001 |
| 4 | 0.89 | 374.85 | 0.01 |
| 5 | 0.82 | 298.04 | 0.16 |
| 6 | 0.80 | 242.21 | 0.45 |
| 7 | 0.77 | 190.98 | 0.78 |
| 8 | 0.71 | 146.27 | 0.95 |
| 9 | 0.68 | 111.48 | 0.99 |
| 10 | 0.67 | 81.32 | 1.00 |
| 11 | 0.59 | 52.26 | 1.00 |
| 12 | 0.49 | 30.90 | 1.00 |
| 13 | 0.46 | 17.36 | 1.00 |
| 14 | 0.27 | 5.32 | 1.00 |
| 15 | 0.17 | 1.51 | 1.00 |

**Table 2.**
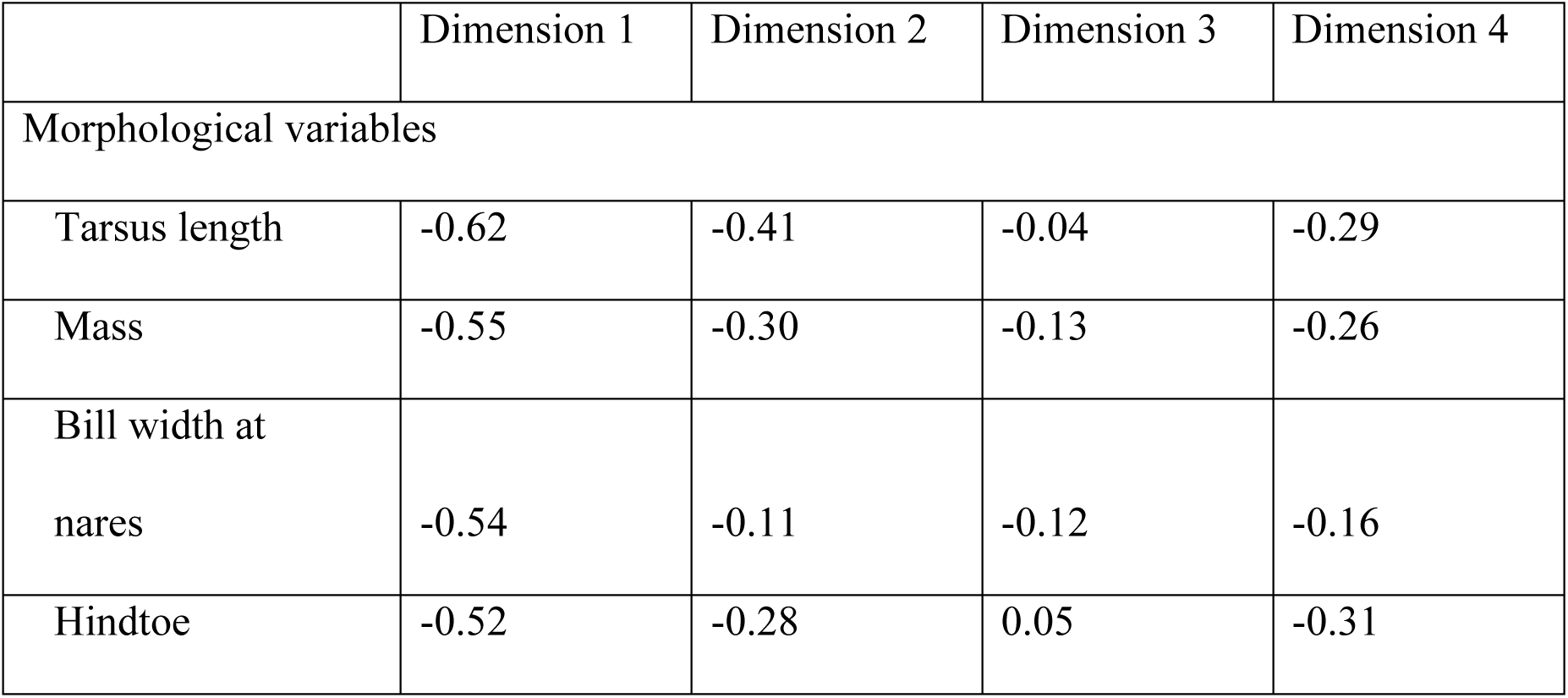
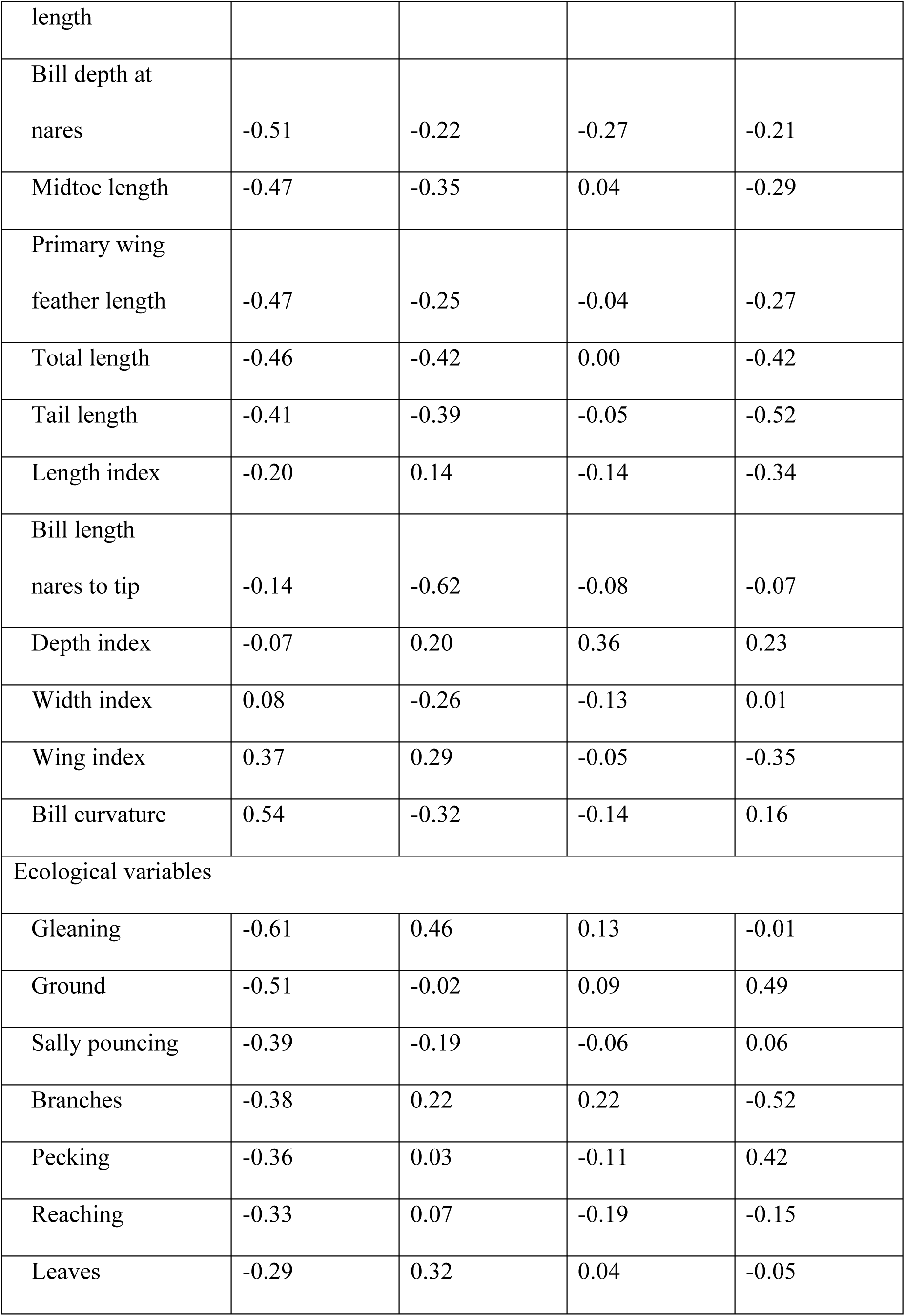
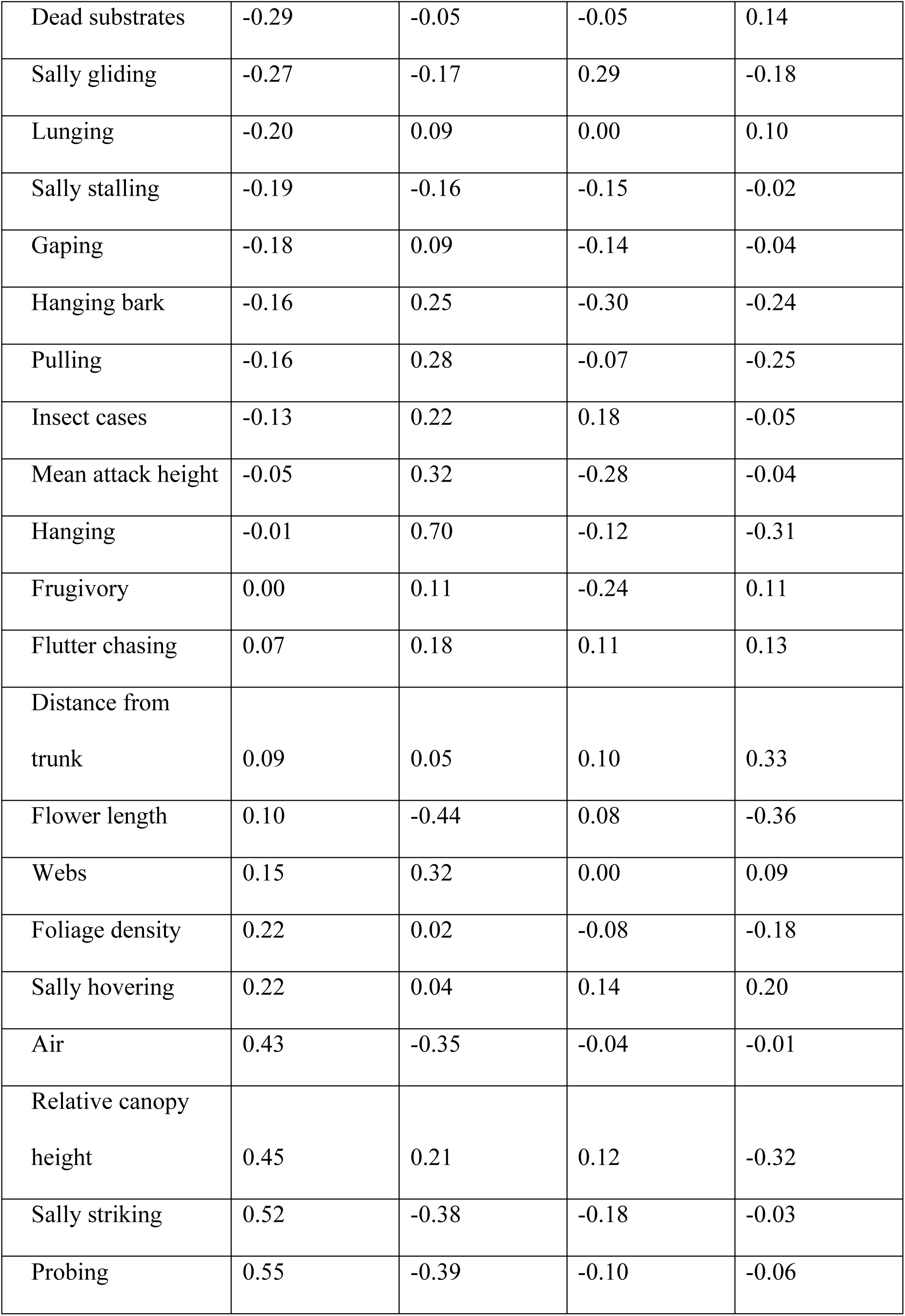
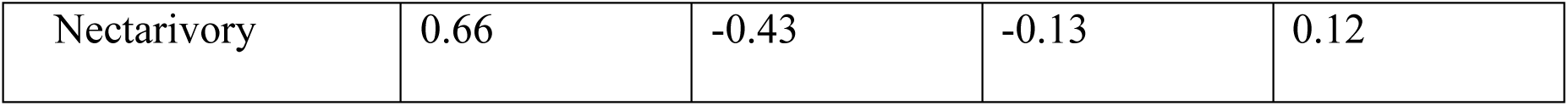
Summary of the phylogenetic correlations of the morphological variables with the morphological canonical correlates (dimensions) and the ecological variables with the ecological canonical correlates (from the phylogenetic canonical correlation analysis).

## Hand-wing index and aspect ratio

The aspect ratio of a wing is the ratio of its span (length) to its breadth (width). In other words, it is a measure of wing shape; higher aspect ratio relates to increasingly long, thin wings. The hand-wing index (Claramunt et al. 2012) is a proxy for aspect ratio, and is a composite of two measures that can be taken on traditionally prepared birds. The appropriate secondary wing feather to measure is the outermost secondary. On our traditionally prepared specimens, we inadvertently measured the length of the longest secondary. Thus, we used our spread wing measures to confirm that the bias from this mistake was small. Across all traditional specimens for which we had corresponding spread wings, the *r^2^* between the longest and outermost secondary wing feathers was 0.99 (*n* = 42). That is because these are generally the same feathers, and we therefore used the measured longest secondary feather to calculate a hand-wing index as in Claramunt et al. (2012).

To confirm that this index was meaningful in Meliphagidae, we calculated aspect ratio using the root box method (Pennycuick 2008). For this, we assumed that the height (perpendicular to the axis of wing) of the root box was equal to the width of the wing (as measured from ImageJ), and that the width of the box (parallel to axis of wing) was equal to the wingspan minus two times the length of the wing. Because this measure relied on a number of assumptions, we derived an alternative measure of aspect ratio as the squared length of the folded primary divided by the wing area (from ImageJ). Both measures were significantly correlated with the hand-wing index (aspect ratio *r^2^* = 0.29, *p* < 0.001, *n* = 64; alternative measure *r^2^* = 0.41, *P* < 0.001, *n* = 69). We therefore used the index as our measure of wing shape, and did not use measures from the spread wing specimens for any additional analyses.

## Additional details on foraging substrate, maneuver, foliage density and distance from the trunk

We recognized 17 mutually exclusive foraging maneuvers (or attacks). In order from most to least frequently observed, these were: probe, glean, sally-strike, sally-hover, flutter-chase, sally-stall, sally-pounce, gape, lunge, peck, sally-glide, flush-pursue, pull, screen, flake, hammer and leap. Definitions of these attacks follow Remsen and Robinson (1990).

We recognized 10 mutually exclusive foraging substrates. In order from most to least frequently observed, these were: flower, leaf, branch, air, ground, fruit, insect case (e.g., Psychidae larvae), web (e.g., picking spiders or their prey out of their webs, or picking caterpillars out of tents), hanging bark, and woody fruit (e.g., *Banksia* cones or *Eucalyptus* capsules).

We recorded foliage density on an ordinal scale, from 0–5. We estimated this density based on a 1-m-diameter sphere centered on the foraging attack site. A zero indicated that 0% of the light passing through that sphere would be blocked, while a five indicated that 100% would be blocked.

We also recorded the distance of the foraging attack from the trunk on an ordinal scale, from 1–4. If the bird was on the trunk we considered it a 1. If the bird was foraging in the middle of the canopy or, for instance, within a hedge of overlapping bushes, we considered it a 2. If the bird was foraging towards the twig tips or just beyond them, for instance hanging from branch tips or sally-striking for insects just beyond the canopy, we considered it a 3. Finally, if the bird was foraging far out in the open, for instance in an open gibber (gravel) expanse or sally-striking for insects far above the canopy, we considered it a 4.

## Introduction to the morphological dataset

The morphological dataset contains measures for 74 of 75 Australian species. We were not able to obtain measures for Eungella Honeyeater (*Bolemoreus hindwoodi*). We included 15 measures in the dataset (Fig. S1–2). (1) Bill length nares to tip is the linear distance in millimeters measured from the distal edge of the nares to the tip of the bill. We also measured bill length from the functional hinge (the base) of the bill to its tip. Thus, (2) length index is defined as:

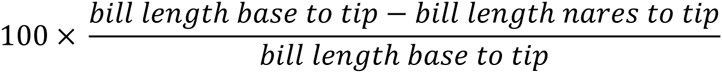

(3) Bill width at the nares is the linear distance in millimeters measured over the under mandible and spanning the distal edge of the nares. (4) The width index is defined as the length index, except that length base to tip and length nares to tip are replaced by width at the base and width at the nares, respectively. (5) Bill depth at the nares is the linear distance spanning the upper and lower mandibles over the distal edge of the nares. (6) Depth index is defined as the width index, except using depth at the base and at the nares of the bill. (7) Bill curvature is defined as the quotient of the length of the upper mandible as measured along its edge in ImageJ over the chord of the bill (also from ImageJ). (8) Primary length is the length in millimeters of the longest primary wing feather as measured with a wing ruler on a traditionally prepared specimen. (9) Tail length is the length in millimeters as measured with a wing ruler pressed up against the crissum of the bird. (10) Tarsus length is the length in millimeters along the tarsus as measured from the tibiotarsal articulation to the base of the toes. (11) Midtoe length is the length in millimeters as measured from the base of the toe to its tip/the start of the nail. Because this toe is often bent in specimens, this was often measured in two or three segments, summing each for a final midtoe length. (12) Hindtoe length is the length in millimeters as measured from the base of the toe to its tip. (13) Total length is the length in millimeters of the entire bird, from the tip of the bill to the tip of the tail. (14) Mass is presented in grams, and is either taken directly from specimens tags or inferred as detailed in the main text. (15) The wing index is calculated like the length index, only instead of bill length from base to tip and from nares to tip, primary and secondary wing lengths, respectively, are used (with the caveat explained in *Hand-wing index and aspect ratio* above).

**Figure S1.**
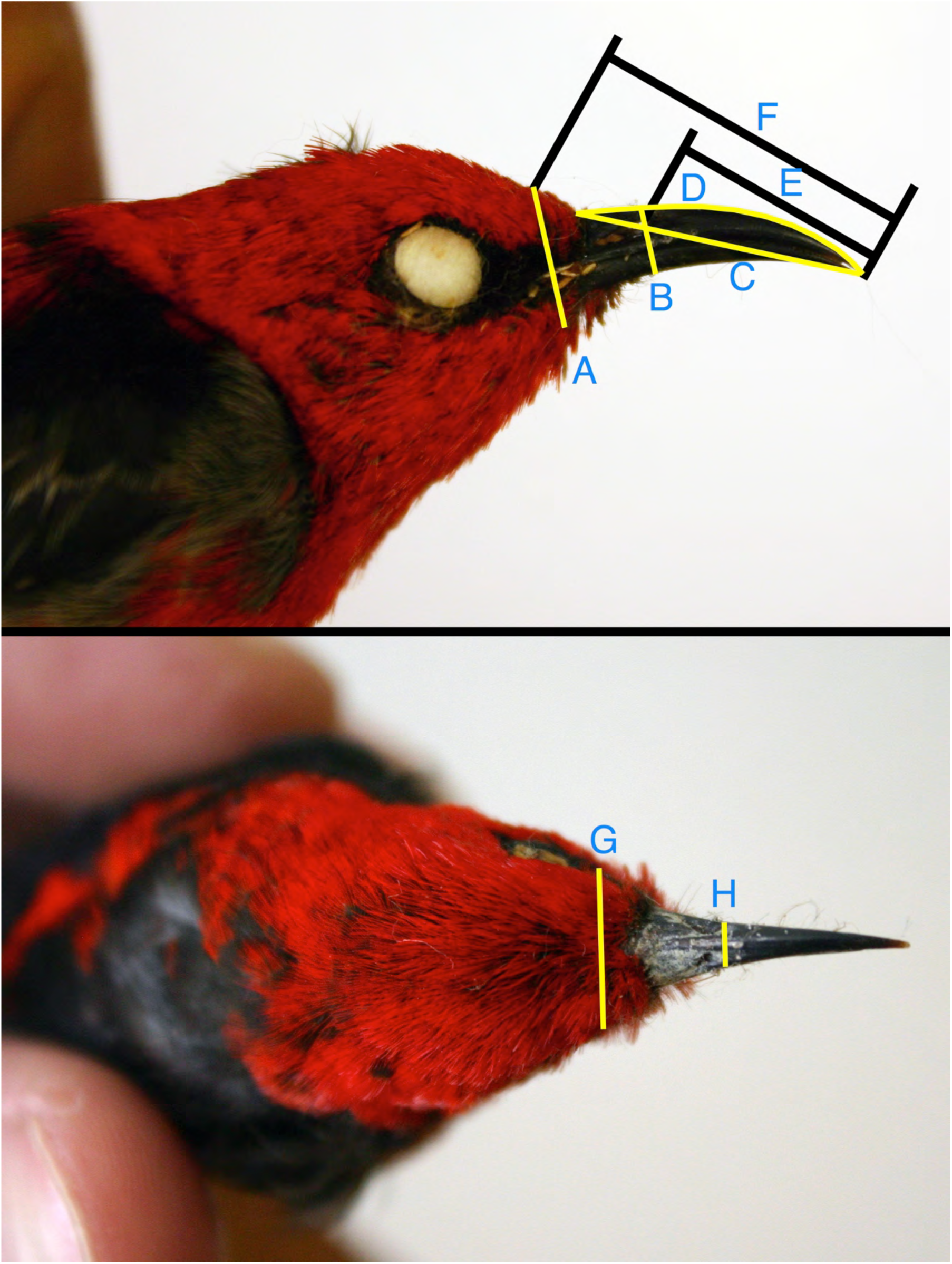
The linear bill measurements recorded in this study, as illustrated on a male Scarlet Myzomela (*Myzomela sanguinolenta*). The measurements are: (A) bill depth at the kinetic hinge, (B) bill depth at the nares, (C) bill chord length, (D) maxilla length, (E) bill length from the nares, (F) bill length from the kinetic hinge, (G) bill width at the kinetic hinge, and (H) bill width at the nares.

**Figure S2.**
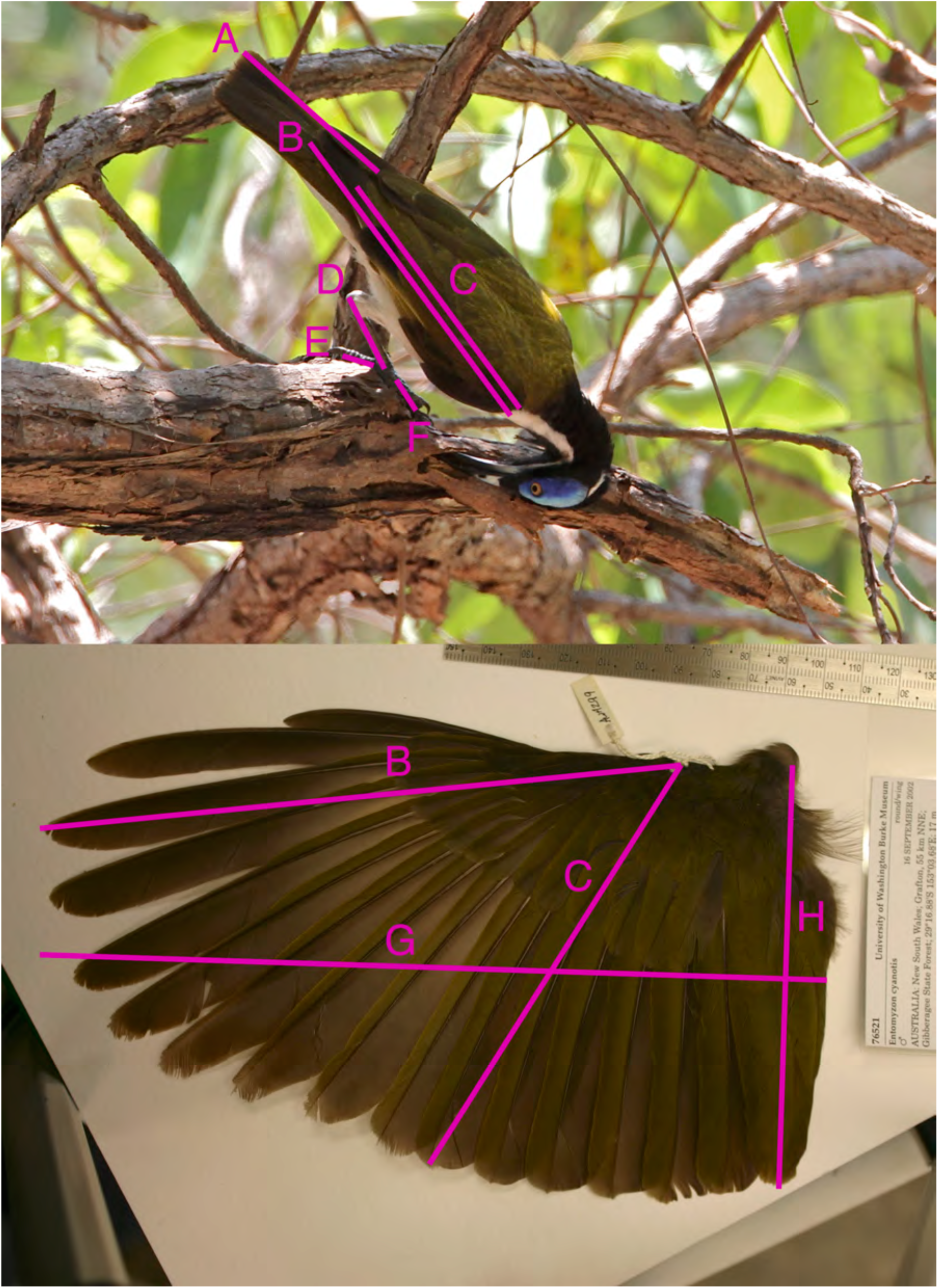
The linear wing, leg and foot measurements taken in this study, as illustrated on a live Blue-faced Honeyeater (*Entomyzon cyanotis*) shown hanging down to probe a dead branch (photo by Bryan Suson) and a prepared Blue-faced Honeyeater spread wing specimen from the Burke Museum. The measurements are: (A) tail length, (B) primary wing length, (C) secondary wing length, (D) tarsus length, (E) hindtoe length, (F) midtoe length, (G) wing length along axis of wing, and (H) wing width perpendicular to axis of wing.

## Introduction to the ecological dataset

The ecological dataset contains measures for 74 of 75 Australian species. We were not able to obtain sufficient observations for Gray Honeyeater (*Conopophila whitei*) to justify inclusion here. The 30 measures included in the main text are presented, as are the number of independent observations per species (i.e., serial observations on a single individual only count as a single observation here). For each foraging observation we recorded the height at which the attack occurred. Average foraging height per species is presented here. For each attack, we also recorded relative attack height. Here, the height of the attack was divided by the average canopy height in a 2 m diameter circle around the attack site; because it was possible for attacks to be above the average canopy height, extreme relative attack height values were rounded down to 1.1. We present the mean relative attack height per species here, where a 1 signifies that species tend to forage at the top of the canopy. We also present the mean foliage density and distance from the trunk, as explained in *Additional details on foraging substrate, maneuver, foliage density and distance from the trunk* above. We include a column for hanging. Hanging is a modifier—attacks like gleaning and probing can be performed while hanging. Thus, the value presented here is the species-specific proportion of observed attacks made while hanging. Reaching, like hanging, is a modifier. It refers to foraging attacks where the legs or neck are completely extended to facilitate the attack. All of the foraging maneuvers (attacks) are explained in detail in Remsen and Robinson (1990). This includes columns 9–20 (columns I through T when opened in Microsoft Excel). Columns 21–30 (U to AD in Excel) refer to the species-specific proportion of attacks that were directed at that substrate. For instance, a species with a 0.5 value for nectarivory was observed foraging on flowers in 50% of the total observations. Dead refers to whether or not the observed substrate was dead. This was most typically tabulated for attacks directed at dead leaves, as frequently employed by *Xanthotis* species. Mean flower length is the average length of the flower fed on by that species. Note that species that were never recorded to forage on flowers are here provisionally coded as a zero; this might best be replaced by NA, depending upon the analysis question.

## Assemblage phylogenetic diversity, trait disparity and climate correlates – unstandardized values and standardized using the richness null model

Our unstandardized measures of phylogenetic distance were non-abundance-weighted (hereafter, “unweighted”) mean pairwise phylogenetic distance (MPD_phylo_) and MPD_phylo_ where the interspecific distances were weighted by species’ abundances in the community in question. Our measures of trait distance were analogous to unweighted and abundance-weighted mean pairwise Euclidean distances in multivariate morphological and ecological space (MPD_trait_). Rather than unstandardized MPD, most of our analyses employed a standardized effect size (SES) metric calculated after randomization of the observed community data matrix with a null model. In the main text we present results as standardized with a null model that maintains species richness, occurrence frequency, and dispersal probability. Here, we present results of assemblage phylogenetic diversity and trait disparity using unstandardized MPD and those standardized with a simpler null model that only maintains species richness (the richness null model of *picante*, Kembel et al. 2010).

Unstandardized MPD_phylo_ was positively correlated with unstandardized morphological MPD_trait_. The same was true whether these MPD values were abundance-weighted (*r*^2^ = 0.40, *p* < 0.001) or not (*r*^2^ = 0.21, *p* < 0.001). That is, assemblages composed of distantly related species were characterized by morphologically disparate species. Unstandardized MPD_phylo_ did not, however, show any clear correlations with unstandardized ecological trait disparity. When not abundance-weighted, there was a very weak negative correlation between the two (*r*^2^ = 0.03*, p* < 0.001). The relationship was not significant when abundance-weighted.

When unweighted morphological MPD_trait_ was standardized with the richness null model, none of the observed communities were considered significantly clustered in morphospace, while three were considered overdispersed. The mean SES was 0.07. When abundance-weighted, no sites were significantly clustered in morphospace, four were overdispersed, and the mean SES was 0.02. From an ecological perspective, using this simple null model and unweighted MPD_trait_, 77 of the communities were considered clustered in ecological space. Ten of the communities were considered overdispersed. The mean SES was −0.49. When abundance-weighted, 72 of the communities were considered clustered, none were considered overdispersed, and the mean SES was −1.27.

There was a weak negative correlation between morphological SES MPD_trait_ and mean annual temperature both when unweighted and abundance-weighted (*r^2^* = 0.05, *p* < 0.001 for both). There was a fairly strong positive correlation between morphological SES MPD_trait_ and mean annual precipitation both when unweighted (*r*^2^ = 0.36, *p* < 0.001) and when abundance-weighted (*r*^2^ = 0.47, *p* < 0.001). Thus, like our preferred dispersal null model, results were qualitatively similar even with a simple richness null model: co-occurring honeyeaters are morphologically more similar in drier and, to a lesser degree, warmer sites.

Ecological SES MPD_trait_ and mean annual temperature were negatively correlated, both when unweighted (*r*^2^ = 0.21*, p* < 0.001) and abundance-weighted (*r*^2^ = 0.13*, p* < 0.001). Morphological SES MPD_trait_ and mean annual precipitation also were negatively correlated, both when unweighted (*r*^2^ = 0.26, *p* < 0.001) and abundance-weighted (*r*^2^ = 0.11, *p* < 0.001). Thus, like our preferred dispersal null model, results were qualitatively similar even with a simple richness null model: co-occurring honeyeaters are ecologically disparate in dry and cold sites.

Honeyeater morphological diversity increased with phylogenetic diversity. Both when unweighted and (*r*^2^ = 0.21, *p* < 0.001) abundance weighted, SES MPD_trait_ was correlated with SES MPD_phylo_ (*r*^2^ = 0.36*, p* < 0.001). Ecological diversity was very weakly negatively correlated with phylogenetic diversity, both when unweighted (*r*^2^ = 0.07, *p* < 0.001) and abundance-weighted (*r*^2^ = 0.01, *p* = 0.01).

## Phylogenetic and trait fields – unstandardized values and standardized using the richness null model

When using a null model that only maintains species richness, species tended to occur in phylogenetically clustered assemblages across their ranges. This is not surprising, since it is essentially the same result as Miller et al. (2013), recapitulated in a species-centered format. As a reminder, a species’ standardized phylogenetic field is a measure of its relatedness to the other species it co-occurs with across its range, standardized to null model expectations. The trend for species to occur in phylogenetically clustered assemblages was not significant when unweighted, where the mean standardized phylogenetic field was −0.50 (Wilcoxon signed rank test that *μ* = 0, *p* = 0.62). When abundance-weighted by species relative abundances, the mean field was −4.61 (Wilcoxon signed rank test *p* = 0.001).

Morphologically, when using the richness null model, species tended to occur in overdispersed trait fields. When unweighted, the mean trait field was 4.18 (Wilcoxon signed rank test *p* < 0.001). When abundance-weighted, the mean was 2.79 (Wilcoxon signed rank test *p* < 0.001).

Ecologically, however, when using the richness null model, species tended to occur in clustered trait fields. This is presumably because species from similar habitats resembled each other in many broad foraging characteristics such as average canopy height and foliage density. Indeed, this was one impetus for the development of the dispersal null model. When unweighted, the mean ecological trait field was −8.36 (Wilcoxon signed rank test *p* < 0.001). When abundance-weighted the mean was −10.23 (Wilcoxon signed rank test *p* < 0.001).

Like the results with the dispersal null model, when standardized with a richness null model there were strong correlations between species’ phylogenetic fields and trait fields. Using phylogenetic generalized least squares regressions, species from phylogenetically overdispersed assemblages tended to occur with morphologically dissimilar species, both when unweighted (*r*^2^ = 0.57, *p* < 0.001) and when abundance-weighted (*r*^2^ = 0.30, *p* < 0.001). This relationship held with unstandardized MPD_phylo_ and MPD_trait_, both when unweighted (*r*^2^ = 0.57, *p* < 0.001) and abundance-weighted (*r*^2^ = 0.40*, p* < 0.001).

As with the dispersal null model, species phylogenetic and ecological trait fields did not exhibit a clear relationship when assessed with a richness null model. The lack of a relationship here is telling—despite notable variation in the phylogenetic neighborhood species find themselves in, there is no consistent pattern in the ecological similarity of these species to those species with which they co-occur. Thus, when standardized phylogenetic and trait field metrics were unweighted, we observed no significant relationship between the two (*r*^2^ = 0.02, *p* = 0.23); abundance-weighting resulted in a weak significant positive relationship (*r*^2^ = 0.08, *p* = 0.02). There was a significant negative correlation when unstandardized, unweighted distances were used (*r*^2^ = 0.35*, p* < 0. 001), but not when unstandardized abundance-weighted distances were used (*r*^2^ = 0.02, *p* = 0.30).

To summarize these results as compared with those in the main text, when dispersal probabilities and species’ occurrence frequencies were not accounted for, many more species were found to occur in phylogenetically clustered assemblages. Moreover, many more species were found to occur in assemblages of ecologically similar species.

Despite these differences, the general pattern of species from phylogenetically clustered assemblages occurring with morphologically similar but ecologically dissimilar species was not obfuscated by using the richness null model. This model makes the potentially unrealistic assumption of allowing any species to occur with equal probability in any assemblage across the continent.

## LITERATURE CITED

Adams, D. C. 2014. A generalized K statistic for estimating phylogenetic signal from shape and other high-dimensional multivariate data. Systematic Biology 63:685–697.

Allen, J. A. 1877. The influence of physical conditions in the genesis of species. Radical Review 1:108–140.

Arnold, S. J. 1992. Constraints on phenotypic evolution. The American Naturalist 140:S85–S107.

Blomberg, S. P., T. Garland Jr, and A. R. Ives. 2003. Testing for phylogenetic signal in comparative data: behavioral traits are more labile. Evolution 717–745.

Bock, W. J., and H. Morioka. 1971. Morphology and evolution of the ectethmoid-mandibular articulation in the Meliphagidae (Aves). Journal of Morphology 135:13–50.

Claramunt, S., E. P. Derryberry, J. V. Remsen, and R. T. Brumfield. 2012. High dispersal ability inhibits speciation in a continental radiation of passerine birds. Proceedings of the Royal Society of London B: Biological Sciences 279:1567–1574.

Cornwell, W. K., D. W. Schwilk, and D. D. Ackerly. 2006. A trait-based test for habitat filtering: Convex hull volume. Ecology 87:1465–1471.

Darwin, C. 1859. On the origin of species by means of natural selection, or the preservation of favoured races in the struggle for life. John Murray, London.

Dehling, D. M., S. A. Fritz, T. Töpfer, M. Päckert, P. Estler, K. Böhning-Gaese, and M. Schleuning. 2014. Functional and phylogenetic diversity and assemblage structure of frugivorous birds along an elevational gradient in the tropical Andes. Ecography 37:1047–1055.

Douglas, M. E., and W. J. Matthews. 1992. Does morphology predict ecology? Hypothesis testing within a freshwater stream fish assemblage. Oikos 213–224.

Fitzpatrick, J. W., J. M. Bates, K. S. Bostwick, I. C. Caballero, B. M. Clock, A. Farnsworth, P. A. Hosner, et al. 2004. Family Tyrannidae (Tyrant-Flycatchers). Pages 170–462 *in*Handbook of the Birds of the World (Vol. 9). Lynx Editions, Barcelona.

Ford, H. A. 1990. Relationships between distribution, abundance and foraging specialization in Australian landbirds. Ornis Scandinavica 133–138.

Ford, H. A., and D. C. Paton. 1976a. Resource partitioning and competition in honeyeaters of the genus Meliphaga. Australian Journal of Ecology 1:281–287.

Ford, H. A., and D. C. Paton. 1976b. The value of insects and nectar to honeyeaters. Emu 76:83–84.

Ford, H. A., and D. C. Paton. 1982. Partitioning of nectar sources in an Australian honeyeater community. Australian Journal of Ecology 7:149–159.

Futuyma, D. J. 2010. Evolutionary constraint and ecological consequences. Evolution 64:1865–1884.

Givnish, T. J. 2015. Adaptive radiation versus “radiation” and “explosive diversification”: why conceptual distinctions are fundamental to understanding evolution. New Phytologist 207:297–303.

Givnish, T. J., and K. J. Sytsma. 2000. Molecular evolution and adaptive radiation. Cambridge University Press, Cambridge.

Gómez, J. P., G. A. Bravo, R. T. Brumfield, J. G. Tello, and C. D. Cadena. 2010. A phylogenetic approach to disentangling the role of competition and habitat filtering in community assembly of Neotropical forest birds. Journal of Animal Ecology 79:1181–1192.

Hansen, T. F., and D. Houle. 2008. Measuring and comparing evolvability and constraint in multivariate characters. Journal of Evolutionary Biology 21:1201–1219.

Higgins, P. J., J. M. Peter, and W. K. Steele. 2001. Handbook of Australian, New Zealand and Antarctic Birds. Vol. 5: Tyrant-flycatchers to Chats. Oxford University Press, Melbourne, Australia.

Joseph, L., A. Toon, Á. S. Nyári, N. W. Longmore, K. Rowe, T. Haryoko, J. Trueman, et al. 2014. A new synthesis of the molecular systematics and biogeography of honeyeaters (Passeriformes: Meliphagidae) highlights biogeographical and ecological complexity of a spectacular avian radiation. Zoologica Scripta 43:235–248.

Karr, J. R., and F. C. James. 1975. Ecomorphological configurations and convergent evolution in species and communities. Pages 258–291 *in* M. L. Cody and J. M. Diamond, eds. Ecology and evolution of communities. Belknap Press, Cambridge, Massachusetts.

Keast, A. 1968. Competitive Interactions and the Evolution of Ecological Niches as Illustrated by the Australian Honeyeater Genus Melithreptus (Meliphagidae). Evolution 22:762–784.

Keast, A., and H. F. Recher. 1997. The adaptive zone of the genus *Gerygone* (Acanthizidae) as shown by morphology and feeding habits. Emu 97:1–17.

Kraft, N. J. B., and D. D. Ackerly. 2010. Functional trait and phylogenetic tests of community assembly across spatial scales in an Amazonian forest. Ecological Monographs 80:401–422.

Lambers, H., and H. Poorter. 2004. Inherent variation in growth rate between higher plants: a search for physiological causes and ecological consequences. Advances in Ecological Research 34:283–362.

Latham, R. E., and R. E. Ricklefs. 1993. Global patterns of tree species richness in moist forests: energy-diversity theory does not account for variation in species richness. Oikos 67:325–333.

Leisler, B., and K. Schulze-Hagen. 2011. The reed warblers: diversity in a uniform bird family. KNNV Uitgeverij, Zeist, Netherlands.

Lovette, I. J., and R. T. Holmes. 1995. Foraging behavior of American Redstarts in breeding and wintering habitats: implications for relative food availability. Condor 782–791.

Luther, D. 2009. The influence of the acoustic community on songs of birds in a neotropical rain forest. Behavioral Ecology 20:864–871.

Mast, A. R., P. M. Olde, R. O. Makinson, E. Jones, A. Kubes, E. T. Miller, and P. H. Weston. 2015. Paraphyly changes understanding of timing and tempo of diversification in subtribe Hakeinae (Proteaceae), a giant Australian plant radiation. American Journal of Botany Online Early.

McGill, B. J., B. J. Enquist, E. Weiher, and M. Westoby. 2006. Rebuilding community ecology from functional traits. Trends in Ecology & Evolution 21:178–185.

Miles, D. B., and R. E. Ricklefs. 1984. The correlation between ecology and morphology in deciduous forest Passerine birds. Ecology 65:1629–1640.

Miles, D. B., R. E. Ricklefs, and J. Travis. 1987. Concordance of ecomorphological relationships in three assemblages of passerine birds. American Naturalist 129:347–364.

Miller, E. T., D. R. Farine, and C. H. Trisos. 2015. Phylogenetic community structure metrics and null models: a review with new methods and software. bioRxiv 025726.

Miller, E. T., and S. K. Wagner. 2014. The ecology of the Australian sandstone *Meliphaga* honeyeater species. Australian Field Ornithology In press.

Miller, E. T., A. E. Zanne, and R. E. Ricklefs. 2013. Niche conservatism constrains Australian honeyeater assemblages in stressful environments. Ecology Letters 16:1186–1194.

Orians, G. H., and A. V. Milewski. 2007. Ecology of Australia: the effects of nutrient-poor soils and intense fires. Biological Reviews 82:393–423.

Pap, P. L., G. Osváth, K. Sándor, O. Vincze, L. Bărbos, A. Marton, R. L. Nudds, et al. 2015. Interspecific variation in the structural properties of flight feathers in birds indicates adaptation to flight requirements and habitat. Functional Ecology 29:746–757.

Parker, S. A. 1973. The tongues of Ephthianura and Ashbyia. Emu 73:19–20.

Paton, D. C. 1980. The importance of manna, honeydew and lerp in the diets of honeyeaters. Emu 80:213–226.

Paton, D. C., and H. A. Ford. 1977. Pollination by birds of native plants in South Australia. Emu 77:73–85.

Pyke, G. H. 1980. The foraging behaviour of Australian honeyeaters: a review and some comparisons with hummingbirds. Australian Journal of Ecology 5:343–369.

Rabosky, D. L. 2014. Automatic Detection of Key Innovations, Rate Shifts, and Diversity-Dependence on Phylogenetic Trees. PLoS ONE 9:e89543.

Rabosky, D. L., M. Grundler, C. Anderson, P. Title, J. J. Shi, J. W. Brown, H. Huang, et al. 2014. BAMMtools: an R package for the analysis of evolutionary dynamics on phylogenetic trees. Methods in Ecology and Evolution 5:701–707.

R Development Core Team. 2011. R: A language and environment for statistical computing. R Foundation for Statistical Computing, Vienna, Austria, http://www.R-project.org.

Recher, H. F. 1971. Sharing of habitat by three congeneric honeyeaters. Emu 71:147–152.

Recher, H. F., and I. J. Abbott. 1970. The possible ecological significance of hawking by honeyeaters and its relation to nectar feeding. Emu 70:90–90.

Recher, H. F., R. T. Holmes, M. Schulz, J. Shields, and R. Kavanagh. 1985. Foraging patterns of breeding birds in eucalypt forest and woodland of southeastern Australia. Australian Journal of Ecology 10:399–419.

Remsen, J. V., and S. K. Robinson. 1990. A classification scheme for foraging behavior of birds in terrestrial habitats. Studies in Avian Biology 13:144–160.

Revell, L. J. 2009. Size-correction and principal components for interspecific comparative studies. Evolution 63:3258–3268.

Revell, L. J. 2012. phytools: an R package for phylogenetic comparative biology (and other things). Methods in Ecology and Evolution 3:217–223.

Revell, L. J., and A. S. Harrison. 2008. PCCA: A program for phylogenetic canonical correlation analysis. Bioinformatics 24:1018–1020.

Ricklefs, R. E. 1996. Morphometry of the digestive tracts of some passerine birds. Condor 98:279–292.

Ricklefs, R. E. 2011. Applying a regional community concept to forest birds of eastern North America. Proceedings of the National Academy of Sciences 108:2300–2305.

Ricklefs, R. E. 2012. Species richness and morphological diversity of passerine birds. Proceedings of the National Academy of Sciences 109:14482–14487.

Ricklefs, R. E., and D. B. Miles. 1994. Ecological and evolutionary inferences from morphology: an ecological perspective. Pages 13–41 *in*Ecological morphology: integrative organismal biology, P.C. Wainwright & S.M. Reilly (eds.). University of Chicago Press, Chicago.

Ricklefs, R. E., and J. Travis. 1980. A morphological approach to the study of avian community organization. Auk 97:321–338.

Rico-Guevara, A., and M. Araya-Salas. 2014. Bills as daggers? A test for sexually dimorphic weapons in a lekking hummingbird. Behavioral Ecology 21–29.

Saunders, M. B., and R. M. R. Barclay. 1992. Ecomorphology of insectivorous bats: a test of predictions using two morphologically similar species. Ecology 73:1335–1345.

Schluter, D. 1996. Adaptive radiation along genetic lines of least resistance. Evolution 50:1766–1774.

Schluter, D. 2000. The ecology of adaptive radiation. Oxford University Press, New York.

Schneider, C. A., W. S. Rasband, and K. W. Eliceiri. 2012. NIH Image to ImageJ: 25 years of image analysis. Nat Meth 9:671–675.

Snow, D. W., and B. K. Snow. 1980. Relationships between hummingbirds and flowers in the Andes of Columbia. Bulletin of the British Museum of Natural History 38:105–139.

Stayton, C. T. 2015. The definition, recognition, and interpretation of convergent evolution, and two new measures for quantifying and assessing the significance of convergence. Evolution.

Stewart, D., and W. Love. 1968. A general canonical correlation index. Psychological Bulletin 70:160–163.

Stiles, F. G. 1981. Geographical Aspects of Bird-Flower Coevolution, with Particular Reference to Central America. Annals of the Missouri Botanical Garden 68:323–351.

Sullivan, B. L., C. L. Wood, M. J. Iliff, R. E. Bonney, D. Fink, and S. Kelling. 2009. eBird: A citizen-based bird observation network in the biological sciences. Biological Conservation 142:2282–2292.

Suryan, R. M., D. B. Irons, and J. Benson. 2000. Prey switching and variable foraging strategies of Black-legged Kittiwakes and the effect on reproductive success. Condor 102:374–384.

Symonds, M. R., and G. J. Tattersall. 2010. Geographical variation in bill size across bird species provides evidence for Allen’s rule. The American Naturalist 176:188–197.

Tilman, D. 1988. Plant strategies and the dynamics and structure of plant communities. Princeton University Press, Princeton, NJ.

Tobias, J. A., R. Planqué, D. L. Cram, and N. Seddon. 2014. Species interactions and the structure of complex communication networks. Proceedings of the National Academy of Sciences 111:1020–1025.

Toon, A., J. M. Hughes, and L. Joseph. 2010. Multilocus analysis of honeyeaters (Aves: Meliphagidae) highlights spatio-temporal heterogeneity in the influence of biogeographic barriers in the Australian monsoonal zone. Molecular Ecology 19:2980–2994.

Villalobos, F., T. F. Rangel, and J. A. F. Diniz-Filho. 2013. Phylogenetic fields of species: cross-species patterns of phylogenetic structure and geographical coexistence. Proceedings of the Royal Society B: Biological Sciences 280:20122570.

Wainwright, P. C. 2007. Functional versus morphological diversity in macroevolution. Annual Review of Ecology, Evolution, and Systematics 38:381–401.

Webb, C. O., D. D. Ackerly, M. A. McPeek, and M. J. Donoghue. 2002. Phylogenies and community ecology. Annual Review of Ecology and Systematics 33:475–505.

Westoby, M., D. S. Falster, A. T. Moles, P. A. Vesk, and I. J. Wright. 2002. Plant ecological strategies: some leading dimensions of variation between species. Annual Review of Ecology, Evolution, and Systematics 33:125–159.

Wiens, J. J., and M. J. Donoghue. 2004. Historical biogeography, ecology and species richness. Trends in Ecology & Evolution 19:639–644.

Willoughby, N. 2005. Comparative ecology, and conservation, of the Melithreptus genus in the Southern Mount Lofty Ranges, South Australia (B.Sc., Hons). University of Adelaide.

Woinarski, J. C. Z., G. Connors, and D. C. Franklin. 2000. Thinking honeyeater: nectar maps for the Northern Territory, Australia. Pacific Conservation Biology 6:61–80.

Wright, I. J., P. B. Reich, M. Westoby, D. D. Ackerly, Z. Baruch, F. Bongers, J. Cavender-Bares, et al. 2004. The worldwide leaf economics spectrum. Nature 428:821–827.

Yanega, G. M., and M. A. Rubega. 2004. Feeding mechanisms: Hummingbird jaw bends to aid insect capture. Nature 428:615–615.

## LITERATURE CITED

Kembel, S. W., P. D. Cowan, M. R. Helmus, W. K. Cornwell, H. Morlon, D. D. Ackerly, S. P. Blomberg, et al. 2010. Picante: R tools for integrating phylogenies and ecology. Bioinformatics 26:1463–1464.

Pennycuick, C. J. 2008. Modelling the flying bird. Academic Press, Burlington, MA.

